# Estrogen-related receptor signaling counters sarcopenia and preserves exercise fitness in naturally aged mice

**DOI:** 10.64898/2026.06.07.730739

**Authors:** DH Sopariwala, A DeBruine, S Poliakova, E Mosa, E Mann, Citu Citu, Zhongming Zhao, A Kumar, VA Narkar

**Author notes:** Corresponding Author: Vihang A. Narkar., Ph.D., Brown Foundation Institute of Molecular Medicine McGovern Medical School, UTHealth, Houston, TX 77030.

## Abstract

**Background:** Estrogen-related receptor gamma (ERRγ) drives an exercise mimicking aerobic gene program in the skeletal muscle that could be beneficial in aging. We have investigated the effect of chronic ERRγ activation on minimizing sarcopenia.

**Methods:** Experiments were performed in muscle specific ERRγ transgenic (TG) mice and wild type (WT) littermates, at young (4-5 months) and old (24-26 months) age. In the skeletal muscle, global gene expression changes, as well as myofiber histological changes in fiber type, size, vascular supply and neuromuscular junction (NMJ), and mitochondrial content were measured. Functional analysis was performed using in vivo muscle contraction assay. Exercise fitness was measured using treadmill sprint and endurance test. Gene and protein expression was measured using QPCR and Westerns, respectively.

**Results:** ERRγ activates a pan-ERR aerobic program in the skeletal muscle to increase expression of 574 genes including ERRα, mitochondrial homeostasis (e.g. Mfn1, Opa1, Drp1, Fis1, and Tfam), vascularization (e.g. Vegfa, Angpt1, Fgf1), and neuromuscular junction (NMJ) (e.g. Nrp1, Aspa, Ptprm, Cxcr4), simultaneously suppressing the expression of atrophy related genes (e.g. Atrogin1, Traf6, Nedd4, Myd88, p21). ERRγ increases mitochondrial content [Mitochondrial area: old TG vs. WT, 2.00 fold; young TG vs. WT, 1.32 fold], oxidative capacity [NADH-TR activity: old TG vs. WT, 1.20 fold; young TG vs. WT, 1.22 fold] and myofiber type [2a: old TG (687±258) vs. WT (252±71); young TG (797±168) vs. WT (440±76); 2x: old TG 1348±87 vs. WT 976±219; young TG 1131±135 vs. WT 936±84; 2b: old TG (798±103) vs. WT (1628±148); young TG (967±133) vs. WT (1623±189)], and capillarity [capillary-to-myofiber ratio: old TG (3.25±0.19) vs. WT (2.41±0.16); young TG (3.41±0.21) vs WT (2.59±0.2)] and [NMJ number [old TG (67±8) vs. WT (40±9); young TG (77±11) vs WT (77±7)], mitigating age-related loss of NMJ and myofiber cross-sectional area [old TG (1570±147µm^2)^ vs. WT (1692.5±208µm^2^) WT; young TG (1828.15±132.8µm^2^) vs. WT (2109.7±296.8µm^2^)]. ERRγ overexpression preserves muscle contractility with aging [Fatigue resistance: 22.72% reduction in force in old vs. young WT; 3.11% reduction in force between old vs. young TG]. Furthermore, ERRγ maintains exercise fitness in old mice [Running: old TG (2964.52±405m) vs. old WT (910.75±6034m); young TG (2232.43±193.64m) vs. young WT (1366.76±60.76m)].

**Conclusions:** ERRγ drives a pan-ERR and counter sarcopenic gene program enhancing oxidative myofiber type, mitochondrial content, vasculature, and NMJ in aging muscle. Consequently, ERRγ minimizes myofiber atrophy, preserves contractility, and improves exercise fitness in old mice. Therefore, ERRs are potential translational targets for combating sarcopenia.

## 1. INTRODUCTION

Sarcopenia is an age-related decline in muscle mass and strength causing skeletal muscle dysfunction [1]. Myofiber atrophy is a key feature of sarcopenia, which is partly caused by protein imbalance due to impaired mTORC1 signaling [2, 3], proteolysis and autophagy [4]. Atrophy is pre-dominantly observed in fast-twitch glycolytic myofibers, whereas slow-twitch oxidative myofibers are less prone [5, 6]. Mitochondrial dysfunction, oxidative stress, mitophagy and NMJ loss additionally contribute to sarcopenia [7, 8]. Aerobic exercise can minimize age-related decline in muscle mass and function [9]. Exercise counters sarcopenia through multiple effects including improved mitochondrial capacity, decreased oxidative stress, muscle vascularization, and NMJ remodeling in the muscle [9]. However, physically active lifestyle is often challenging due to personal choices or medical issues. Molecular pathways mediating benefits of aerobic exercise in the muscle could be advanced as clinical targets for managing sarcopenia.

Estrogen-related receptors (ERRs) have a major regulatory role in the skeletal muscle [10]. Muscles highly express ERRα and ERRγ, with minimal ERRβ expression [11]. Exercise induces ERRα and ERRγ in the muscle [12]. The impact of ERRs on muscle homeostasis is demonstrated through transgenic studies. Muscle-specific ERRγ overexpression increases mitochondrial biogenesis, fatty acid oxidation and vascularization [11]. Muscle-specific ERRα overexpression drives metabolic and vascular remodeling similar to ERRγ [13]. Individual muscle-specific loss of ERRs have mild effect on the skeletal muscle, whereas compound deletion of ERRs in muscle causes severe mitochondrial deficit, vascular regression, and exercise intolerance [12, 14, 15]. ERRs are mitigative in various myopathy models including Duchenne muscular dystrophy (DMD) [16, 17], peripheral arterial disease (PAD) [18] and ACL injury [19]. While there is a dearth of clinical or human studies on ERRs, there is clear indication of translational potential. Expression of ERRs are increased in muscle biopsies from exercise-trained humans [20, 21], whereas muscle expression is decreased in maladies such as diabetes, sarcopenia and muscular dystrophies [22–24].

Potential beneficial effects of ERR-induced transcriptional program in aging and mitigation of sarcopenia has not been studied. Here, we have investigated the effect of life-long skeletal muscle-specific ERRγ overexpression on sarcopenia and exercise fitness in aged mice. We show that ERRγ drives an ‘anti-aging’ gene program in the muscle to prevent the decline in muscle size, contractile function and exercise fitness with aging.

## 2. METHODS

### 2.1 Experimental overview

We previously described the muscle-specific ERRγ transgenic mice (TG) [11]. Animals were maintained and treated in accordance with the U.S. National Institute of Health Guide for Care and Use of Laboratory Animals. The procedures were approved by the Animal Welfare Committee at The University of Texas Health Science Center in Houston (UTHealth).

All experimental parameters were measured in young (5 months) and old (24-26 months) male TG and littermate wild type (WT) mice. Gene expression, protein expression, histological analysis, and electron microscopy were performed in the gastrocnemius and/or tibialis anterior (TA) muscle. RNA-sequencing and data analysis were performed in gastrocnemius of the TG and WT mice. Physiological testing including treadmill sprinting and endurance test, and in vivo hindlimb muscle functional analysis was performed.

Detailed methodology is described in SUPPLEMENTAL METHODS.

### 2.2 Statistics

Unpaired Student’s t-test and one-way ANOVA with post-hoc test were used, as described in figure legends. Data in figures is presented as mean ± standard error of mean, such that N used in each experiment was sufficient to achieve a power of 0.8 and Type 1 error rate of 5%. Data were analyzed and plotted using GraphPad Prism 9. The analysis of RNA-seq data is described in Supplemental methods.

## 3. RESULTS

### 3.1 Skeletal muscle ERRγ expression in young and old mice

Young (4-5 months) and old (24-26 months) TG mice and their WT littermates were used. We first analyzed *Esrrg* mRNA expression in the gastrocnemius of WT and TG mice, at young and old age. Gastrocnemius from TG mice expressed induced levels of Esrrg mRNA compared to WT mice in both young and old mice (Figure S1a), showing that muscle transgene expression is sustained with aging. Endogenous Esrrg mRNA expression was also sustained in WT muscles with aging. Endogenous Esrra mRNA (another ERR isoform expressed in muscle) was induced in TG vs. WT mice, at both young and old age (Figure S1b). ERRγ and ERRα protein expression was also elevated in gastrocnemius of TG vs. WT mice independent of age (Figure S1c-d). Age-related decline in protein expression of ERRs was not detected. ERRγ was selectively overexpressed in the skeletal muscle (as driven by a muscle-specific promoter). Other tissues (e.g. kidney, liver, adipose, heart) showed comparable ERRγ expression between WT and TG mice (Figure S1e). Next, we measured the effects of muscle-specific ERRγ overexpression on gene expression, and key parameters of muscle homeostasis including oxidative capacity, vascular supply, neuromuscular junction (NMJ), myofiber (type, number, and size), and muscle function in young and old mice.

### 3.2 ERRγ transcriptional program in skeletal muscle

We measured the transcriptomic effect of ERRγ overexpression in the skeletal muscle in young mice, followed by validation of major ERRγ-activated pathways in young and old mice. RNA sequencing in young WT and TG gastrocnemius showed that there were ∼1,100 differentially expressed genes with comparable distribution of up-regulated and down-regulated genes (Figure 1a-b). Majority of the top 20 up-regulated pathways were related to mitochondrial respiration and oxidative metabolism (Figure 1c). On the other hand, the top 20 down-regulated pathways were linked to shift in fiber type and related processes (e.g. calcium ion transport) (Figure 1d).

**Fig. 1.**
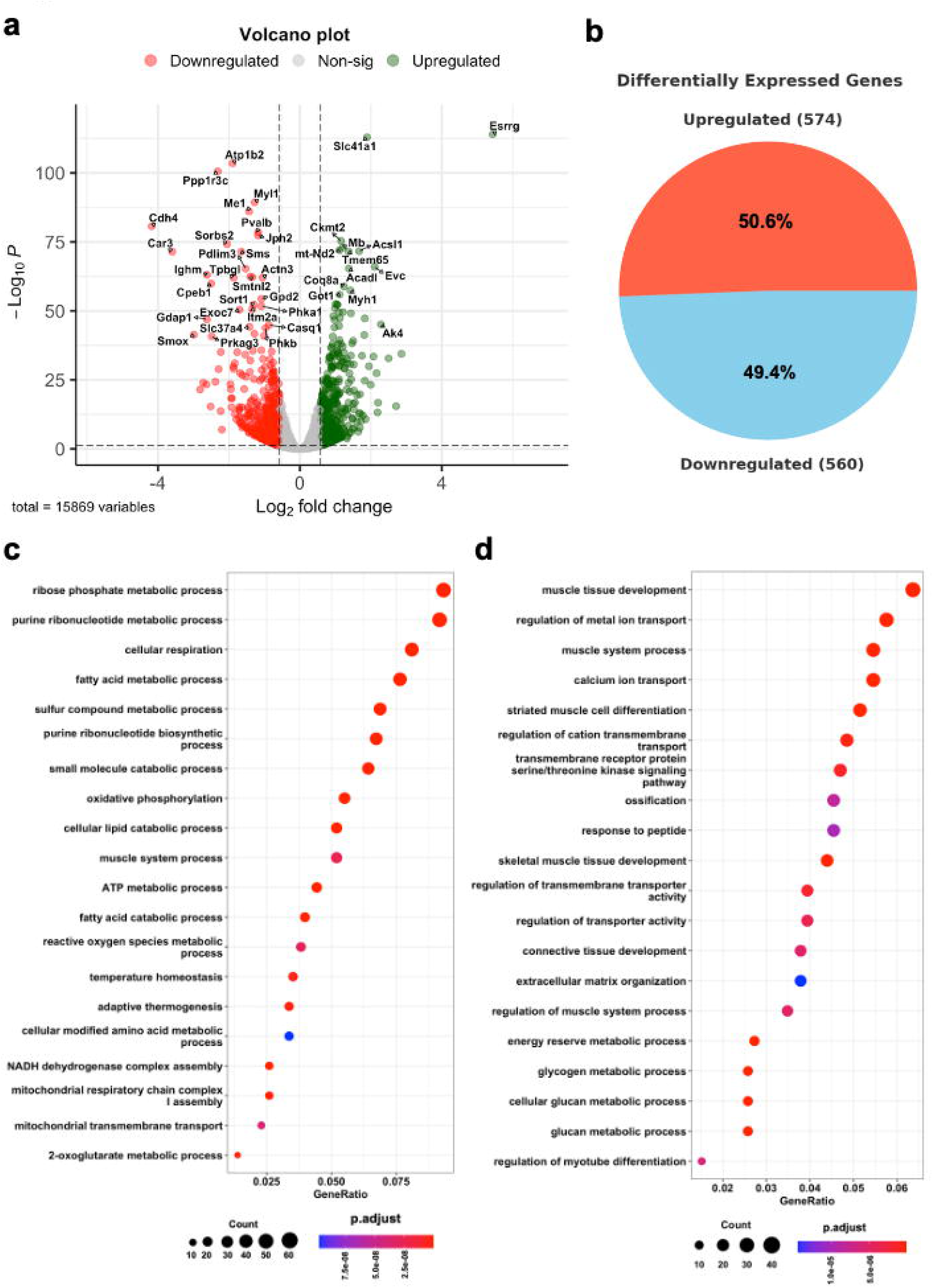
Transcriptomic effects of ERRγ in skeletal muscle. **(a)** Volcano plot showing the distribution of up or down-regulated genes in gastrocnemius from young TG relative to young WT mice (N=3 mice). **(b)** Overexpression of ERRγ in gastrocnemius muscle of TG mice results in similar percentage of differentially expressed genes being up-regulated and down-regulated as compared to WT muscle. **(c-d)** Top 20 up-regulated (c) and down-regulated (d) gene pathways in TG vs. WT gastrocnemius.

### 3.3 ERRγ increases oxidative capacity in young and old muscle

RNA sequencing in young WT and TG mice showed higher expression of mitochondrial genes in TG vs. WT muscle (Figure 2a). We validated key genes associated with mitochondrial biogenesis and dynamics (fission and fusion), and antioxidant defense in muscles of young and old mice. Expression of mitochondrial biogenesis and metabolism (Tfam, Cycs, Cox6a, and Sdhb) and reactive oxygen species (ROS) sequestration (Catalase and Sod2) genes was significantly higher in gastrocnemius of TG vs. WT mice, in both young and old mice (Figure 2b-c). Notably, transcripts trended towards lower expression in old vs. young muscle. Genes involved in mitochondrial fusion (Mfn1 and Opa1) and fission (Drp1 and Fis1) were comparably expressed in TG vs. WT muscles of young mice but were induced in old TG mice (Figure 2d). We measured the protein expression of CYCS and representative subunits from oxidative respiratory chain (Figure S2a) in gastrocnemius. While protein expression was increased in TG vs. WT muscle, we did not detect an age-based difference (Figure S2a). NADH-TR activity staining, indicative of Complex 1 function of mitochondrial electron transport chain, was significantly higher (blue coloration) in TG vs. WT muscles, without any age-dependent differences (Figure S2b).

**Fig. 2.**
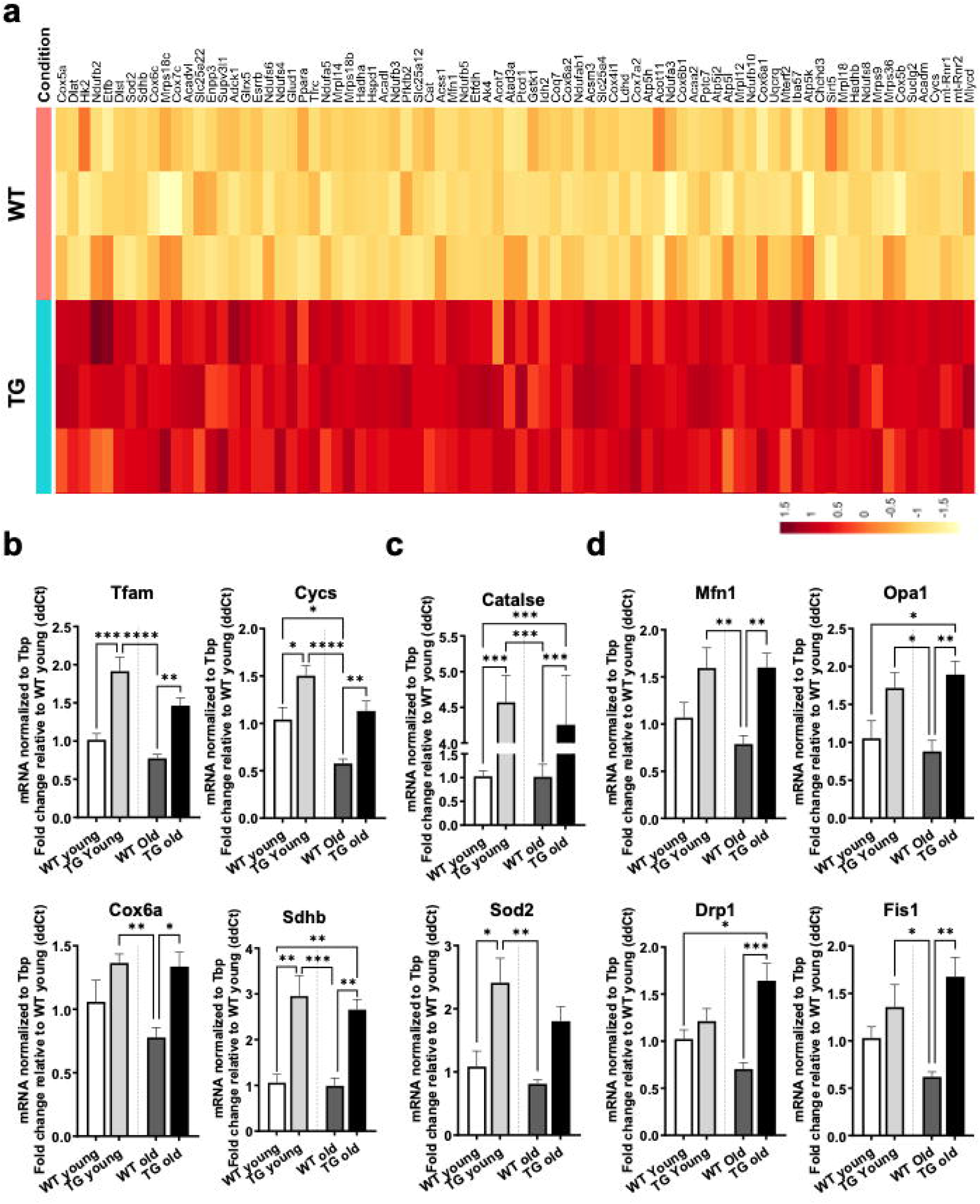
Sustainable induction of oxidative metabolic gene program by ERRγ with aging. **(a)** Heat map showing expression of mitochondrial and oxidative metabolism related genes in TG vs. WT gastrocnemius from young mice (N=3). **(b-d)** Expression of mitochondrial biogenesis (Tfam, Cycs, Cox6a, Sdhb) (b), reactive oxygen species sequestration (Catalase, Sod2) (c), mitochondrial fusion (Mfn1 and Opa1) and fission (Drp1 and Fis1) (d) genes (N=6). One-way ANOVA with Tukey’s post hoc test. p<0.05=*, p<0.01=**, p<0.001=***, and p<0.0001=****.

For in-depth mitochondrial analysis, we performed transmission electron microscopy (TEM) in WT and TG tibialis anterior (TA) muscle to quantify mitochondrial content to validate the increase in mitochondrial oxidative capacity in TG mice. Analysis of intermyofibrillar mitochondria in TG vs. WT TA from young and old mice (Figure 3a) showed a significantly higher total and average mitochondrial area in TG vs. WT TA (Figure 3b-c). Strikingly, old vs. young WT TA had significantly lower total mitochondrial area, however this mitochondrial deficiency was mitigated in old vs. young TG TA (Figure 3b). Mitochondrial number and average area showed significantly lower levels in old vs. young WT TA by t-test but not one-way ANOVA (Figure S3a-c). Similarly, analysis of subsarcolemmal mitochondria (Figure 3d) showed significantly higher total mitochondrial area (Figure 3e) and number (Figure 3f) in TG vs. WT TA of young and old mice. The total mitochondrial area of old vs. young WT TA was lower when analyzed by t-test (Figure S3d). The average mitochondrial area did not show significant difference between the groups (Figure S3e-f). TEM demonstrated a decrease in several morphological mitochondrial homeostatic parameters in old vs. young TA. ERRγ through induction of mitochondrial biogenesis and dynamics genes increases mitochondrial content and oxidative capacity in muscle that prevents mitochondrial decline with aging.

**Fig. 3.**
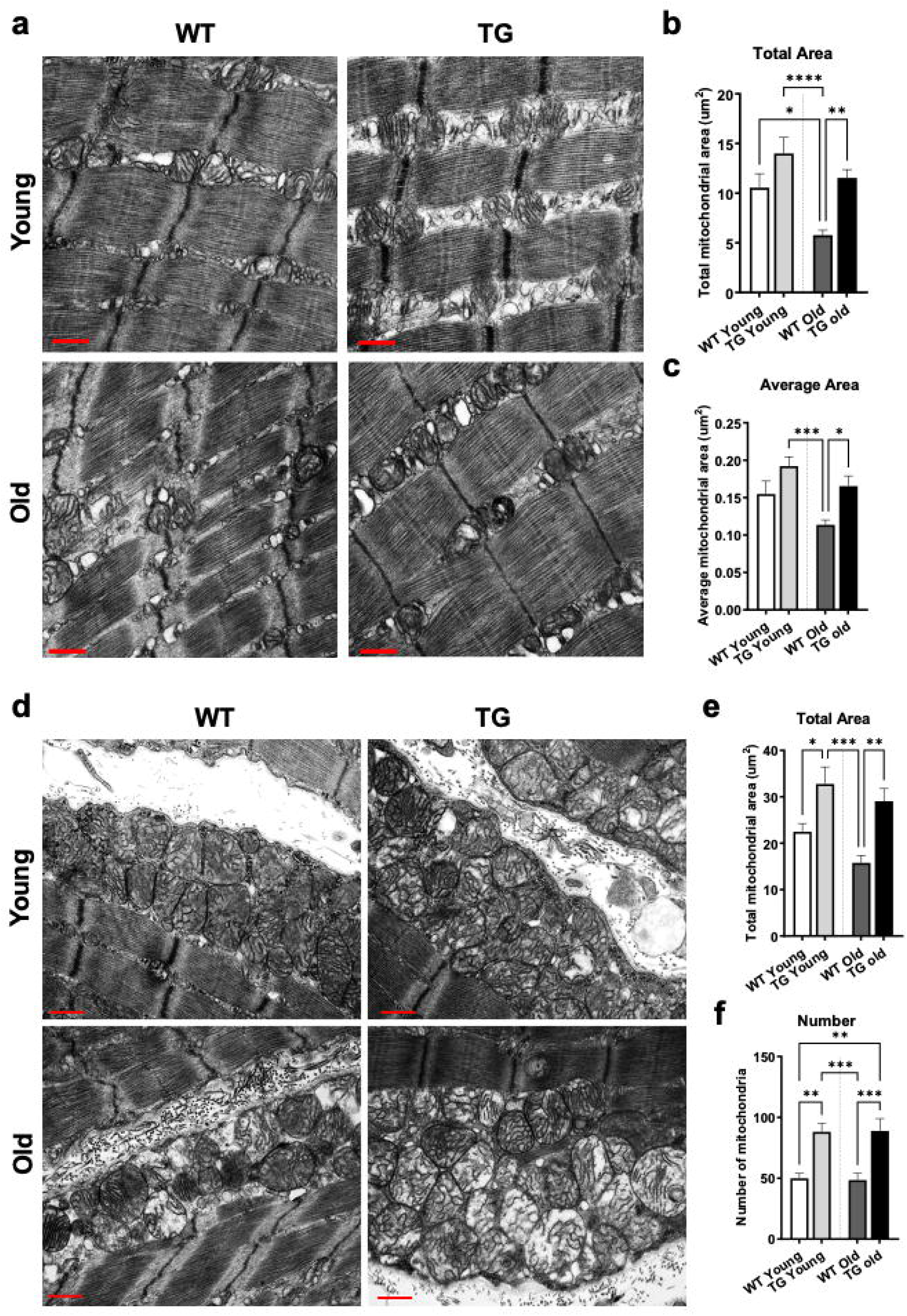
ERRγ prevents age-related decline in mitochondrial content. Following measurements were made in tibialis anterior TA of WT and TG mice at young and old age. **(a-c)** Representative transmission electron microscopy (TEM) images of intermyofibrillar region of tibialis anterior (TA) muscles (a), showing quantification of total mitochondrial area per micrograph (b) and average mitochondrial area (c). **(d-f)** Representative transmission electron microscopy (TEM) images of subsarcolemmal region of TA (d), showing quantification of total mitochondrial area per micrograph (e) and mitochondrial number (f). Red bar = 600 nm. N=20 to 24 micrographs from 3 mice per group. One-way ANOVA with Tukey’s post hoc test. p<0.05=*, p<0.01=**, p<0.001=***, and p<0.0001=****.

### 3.4 ERRγ increases capillary density and neuromuscular junction (NMJ) number in young and old muscle

RNA sequencing in young TG vs. WT mice showed induction of angiogenic genes by ERRγ in the TG gastrocnemius (Figure 4a). ERRγ overexpression in the muscle induced capillary density in the TA in young and old mice. Gastrocnemius in the TG mice have significantly higher protein expression of CD31 (endothelial cell marker) and PDGFRβ (pericyte marker) compared to WT mice in both young and old mice (Figure 4b). Capillary-to-myofiber ratio is significantly higher in TG vs. WT TA at young and old age (Figure 4c). We did not detect an age-related regression of muscle capillarity in mice, at the level of CD31 staining.

**Fig. 4.**
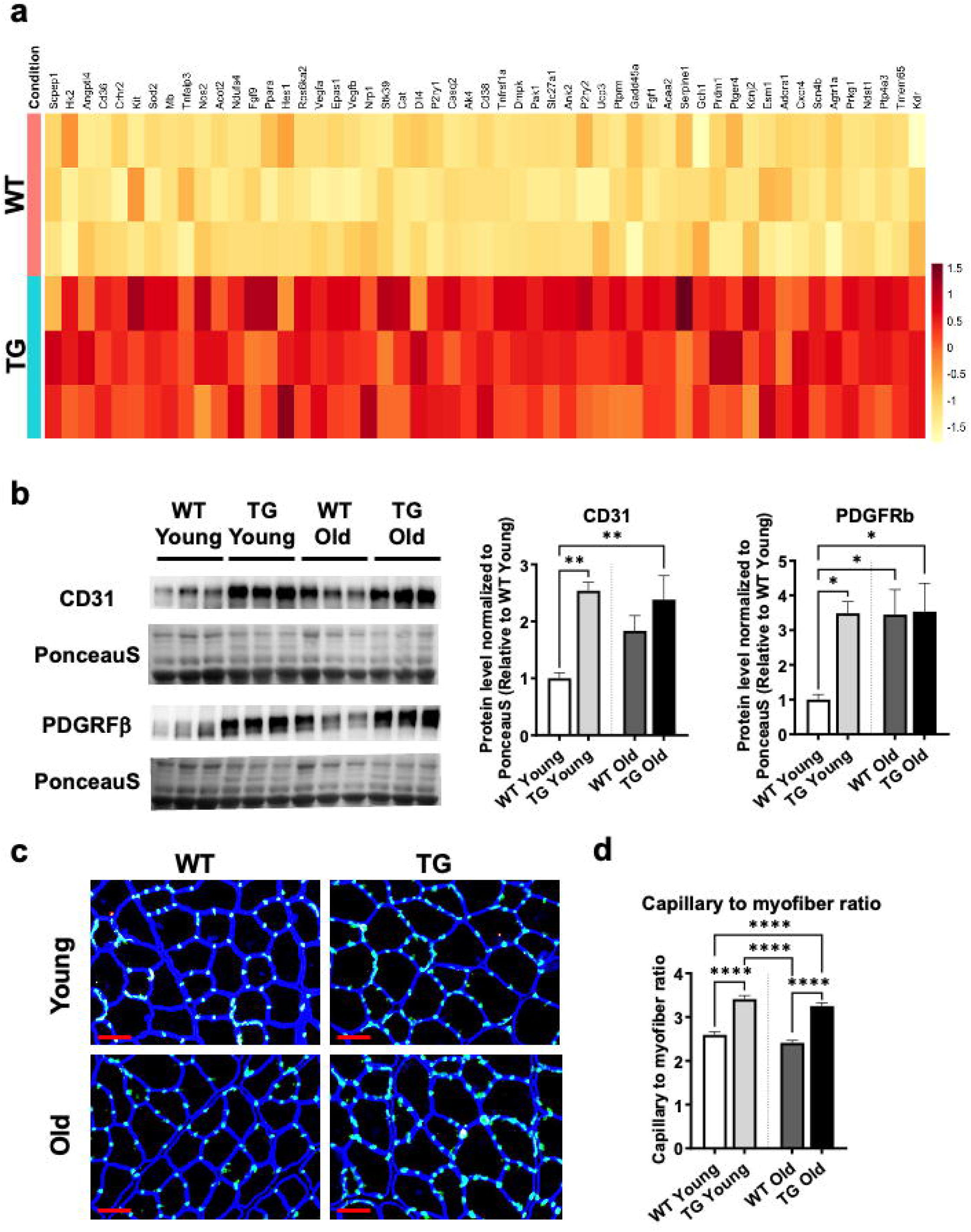
Sustainable induction of vascular gene program by ERRγ in aging muscle. **(a)** Heat map showing expression of genes related to angiogenesis in TG vs. WT gastrocnemius from young mice (N=3). **(b)** Representative western blot and quantification showing expression of CD31 and PDGFRβ proteins in gastrocnemius muscle of young and old TG vs. WT mice (N=6). **(c)** Representative immunostaining of Tibialis anterior (TA) for CD31 (Green, endothelial cells) and laminin (blue, myofiber membrane), and quantification showing capillary to myofiber ratio in TG vs WT muscles from young and old mice (N=5). Red bar = 50µm. One-way ANOVA with Tukey’s post hoc test. p<0.05=*, p<0.01=**, and p<0.0001=****.

Sarcopenia is linked to loss of NMJs [25]. Degradation of NMJ causes ineffective muscle contraction leading to atrophy and weakness [26, 27]. RNA sequencing revealed higher expression of NMJ genes in TG vs. WT gastrocnemius in young mice (Figure 5a). QPCR confirmed induction of genes related to axon guidance and development (Aspa, Nrp1, Ptpm, Cxcr4, and Pcdh17) in TG vs. WT muscle in both young and old mice (Figure 5b). To assess NMJ number in TG vs. WT TA in both the young and old mice, we performed immunostaining for neurofilament (nerve body), synapsin-1 (synaptic junction), and α-Bungerotoxin (acetylcholine receptor) to image overlap between these structures (purple region) (Figure 5c). TG vs. WT TA had elevated NMJ number (purple regions showing overlap between acetyl choline receptor and neurofilament) in both the young and the old mice (Figure 5d). There was a significant decrease in NMJ number in old vs. young WT TA. Notably, the NMJ number were maintained in the old vs. young TG TA (Figure 5e). These data indicate that ERRγ overexpression in the muscle not only increases the number of NMJ in young mice but also prevents the loss of NMJ with aging.

**Fig. 5.**
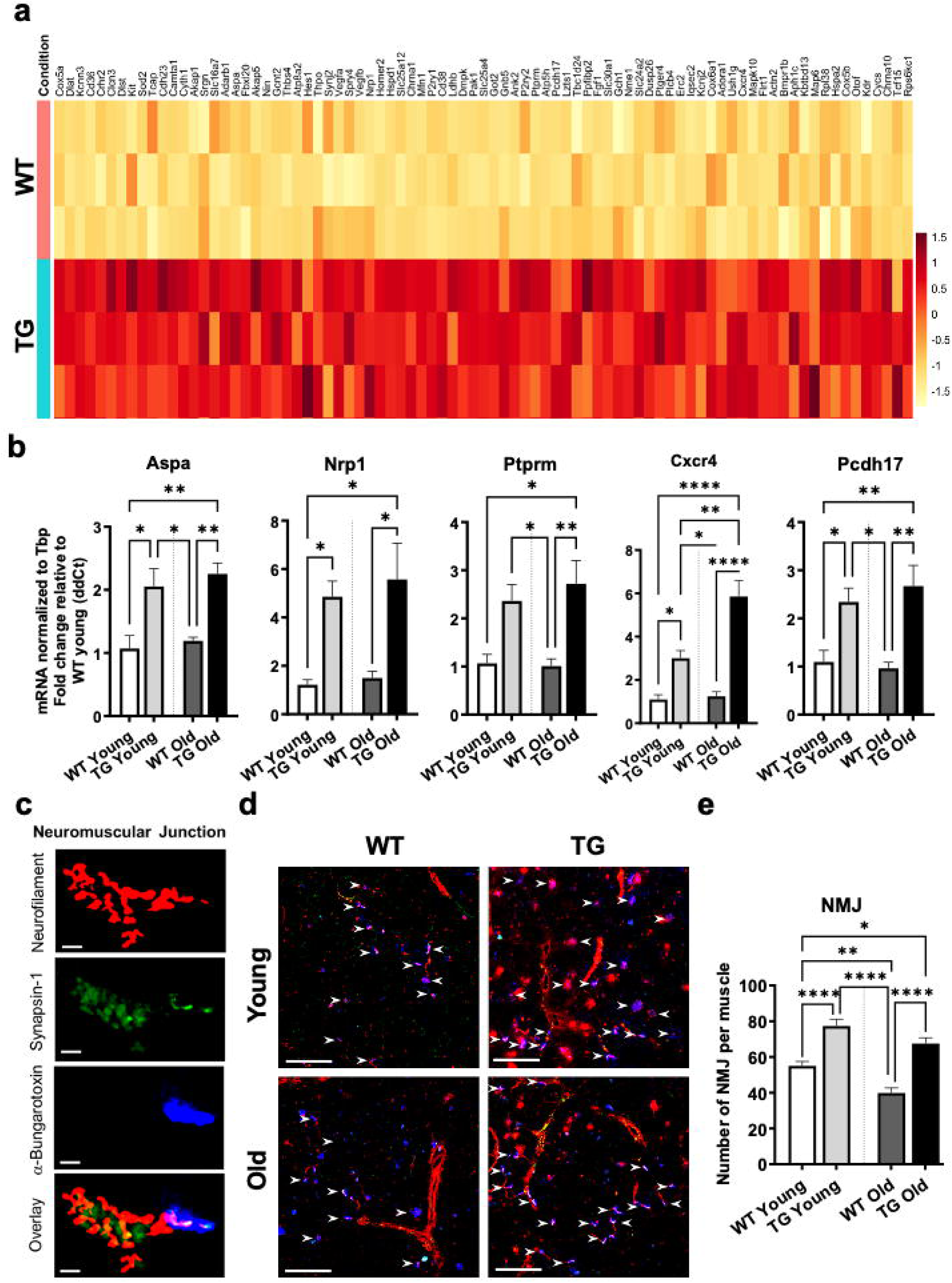
Regulation of neuromuscular junction (NMJ) by ERRγ. **(a)** Heat map showing expression of genes related to NMJ in TG vs. WT gastrocnemius from young mice. (N=3). **(b)** Expression of genes related to axon guidance in TG vs. young gastrocnemius from young and old mice (N=5). **(c)** Representative image showing immunostaining of NMJ components (magenta) in the overlay tile showing a single NMJ. White bar = 20 µm. **(d)** Representative immunostaining images of TG vs. WT TA cryosections from young and old mice (N=5). White arrows indicate NMJ (Magenta). White bar = 200 µm. **(e)** Quantification of the number of NMJ in TG vs. WT muscles from young and old mice (N=5). One-way ANOVA with Tukey’s post hoc test. p<0.05=*, p<0.01=**, and p<0.0001=****.

### 3.5 ERRγ promotes myofiber type remodeling and protects against muscle atrophy

Myofiber type, number, and size analysis in the TA of WT and ERRγ TG mice was performed at young and old age. TA were stained for myosin heavy chain (MyHC) type 2a (oxidative fast), 2x (oxidative/glycolytic fast) and 2b (glycolytic fast) myofibers (Figure S4a). A myofiber type distribution analysis of TG vs. WT TA from young and old mice showed that the total myofiber number between the 4 groups was similar (Figure S4b). Both young and old TG TA had significantly higher number and percentage of type 2a myofibers compared to WT (Figure S4c and S4f). Similarly, type 2x myofiber proportion was significantly higher in both young and old TG vs. WT TA (Figure S4d and S4g). In contrast, type 2b myofiber proportion (number and percentage) was lower in TG vs. WT TA, in both age groups (Figure S4e and S4h). Therefore, ERRγ drives a robust oxidative myofiber switch in young mice, which is sustained with aging in the muscle.

We next measured the effect of muscle ERRγ overexpression on myofiber size (cross-section area) in young and old mice. TA cryosection immunostained for the three major MyHC isoforms (Figure 6a) were quantified for size distribution of each myofiber type. The size distribution curve for total myofibers and each individual myofiber type were constructed to provide insights into the effects of aging and ERRγ overexpression. Type 2b myofiber atrophy (immunostained red) is qualitatively detected in old WT group vs. all other groups (Figure 6a). This is underscored in size distribution curve of type 2b myofibers with the significant leftward shift (smaller myofiber cross-sectional area) in TA of old WT group vs. all other groups (Figure 6b – blue line is old WT TA). Size distribution for myofiber type 2a shows no difference between groups (data not shown). Type 2x myofibers show leftward shift in old vs. young TG TA (Figure 6c). The total myofiber size distribution (Figure 6d) revealed that TA from the old TG mice show leftward shift predominantly in myofibers less than 2000µm^2^, while TA from the old WT mice show leftward shift in larger myofibers with area greater than 2000µm^2^ as compared to young TG and WT TA, respectively.

**Fig. 6.**
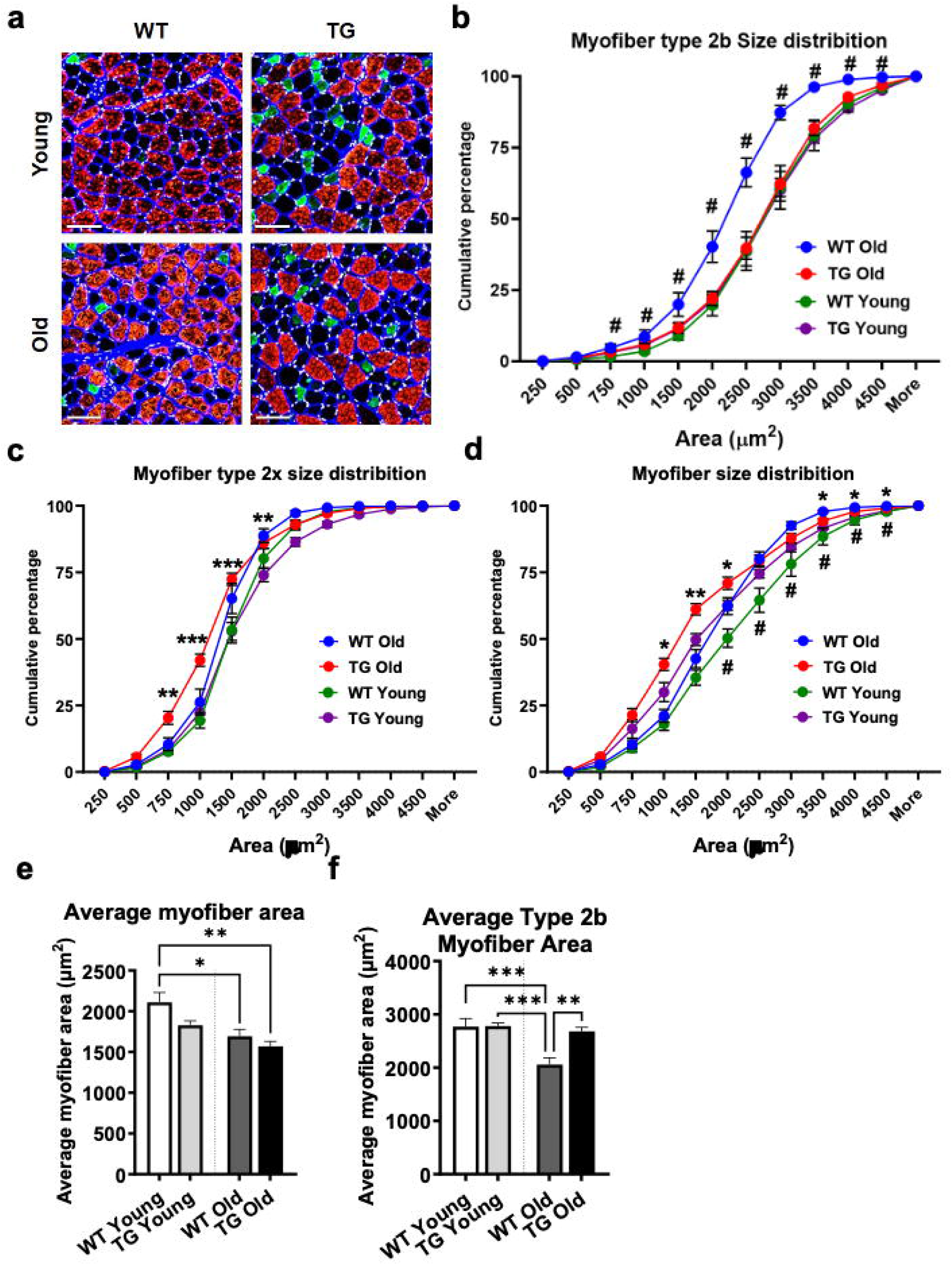
ERRγ regulates myofiber size distribution. Following measurements were made in tibialis anterior (TA) muscle cryosection from WT and TG mice at young and old age. **(a)** Representative image of MHC immunostained TA cross-sections. Red – type 2b, Green – type 2a, Black – type 2x. White bar = 100 µm. **(b)** Myofiber type 2b size distribution. **(c)** Type 2x myofiber size distribution of WT and TG TA from young and old mice. **(d)** Total myofiber size distribution of WT and TG TA from young and old mice TA showing leftward shift of old WT TA relative to young WT TA (#) and leftward shift of old TG TA relative to young TG TA (*). **(e)** Quantification of average myofiber area of all fiber types combined. **(f)** Quantification of average myofiber area of type 2b myofibers (N=5). Unpaired Student’s t-test. p<0.05=*, p<0.01=**, p<0.001=***. P<0.01= #. One-way ANOVA with Tukey’s post hoc test. p<0.05=*, p<0.01=**, and p<0.001=***.

Average myofiber cross-section area and total myofiber cross-section area (product of average area and total myofiber number) was significantly lower in the TA of old vs. young WT mice (Figure 6e and S5c). The age-dependent decline in myofiber size was negated in TA from TG mice (Figure 6e and S5c), comparing old vs. young TG mice. In terms of individual myofiber type, average cross-sectional area of type 2a myofibers were comparable between young and old TA in both the WT and TG groups (Figure S5a), suggesting that the type 2a myofibers are less prone to atrophy. Note that TA of TG vs. WT mice (both young and old) had significantly higher total type 2a myofiber area (Figure S5d) due to the high type 2a myofiber number in TG TA.

The average cross-sectional area of type 2b myofibers is significantly lower in the WT TA from the old vs. young mice. This decline in type 2b average myofiber cross-sectional area was absent in the TG TA, comparing old vs. young TG mice (Figure 6f). Note, TA from the old TG vs. WT mice has significantly higher average cross-sectional area of type 2b myofibers (Figure 6f), indicating that the type 2b myofibers are protected against muscle wasting by ERRγ overexpression in the TG mice. Similarly, the total type 2b myofiber area of TA is significantly lower in old vs. young WT mice (Figure S5e); whereas TA from old vs. young TG mice had comparable total type 2b myofiber area (Figure S5e). Note that the total type 2b myofiber area of TA in young and old TG mice is significantly lower than in age matched WT mice TA (Figure S5e). This is due to the myofiber type switch in TA of the TG vs. WT mice (Figure S4).

Old WT and TG TA have lower average type 2x myofiber area compared to young WT and TG mice (Figure S5b). TA from both old WT and TG mice trending towards lower total type 2x myofiber area vs. young WT and TG mice, without reaching significance (Figure S5b). Although TA from old TG mice have significantly lower average type 2x area than young TA, it is not reflected in the total type 2x myofiber area (Figure S5b and Figure S5f), because of the higher number of type 2x myofibers in the old TG muscles (Figure S4d).

We performed qPCR analysis on key atrophy-related genes (Atrogin1, Nedd4, Traf6, Myd88) and the senescence marker p21 in the gastrocnemius from the four groups (Figure 7a-b). While the expression of these genes is significantly increased in the muscle of the old vs. young WT mice, it was not induced in the muscles of old vs young TG mice (Figure 7a-b). We quantified the centrally located nuclei within the myofibers across four groups, which is an indicator of myofiber turnover [23]. TA from the young TG and WT mice have a low percentage of central nuclei (Figure 7c – yellow arrows point to the white central nuclei within the myofibers). In contrast, the TA of old WT mice had a significantly higher percentage of central nuclei compared to young WT mice. Notably, the percentage of central nuclei was lower in the old TG vs. old WT TA (Figure 7d), indicating suppression of age-dependent muscle damage by ERRγ overexpression. Myogenic factor expression (MyoD, MyoG) was not affected in any of the groups (data not shown). TEM analysis also showed centrally located nuclei along with multivesicular bodies (MVB) in the WT old mice (Figure 7e). As reported [28, 29], we found large areas in the muscle of old WT mice that have aggregates because of SR misfolding (Figure 7f left and middle panel at two different magnifications), within which there were trapped mitochondria (Figure 7f, middle panel: mitochondria are labelled with M). This is a hallmark of arcopenia [28, 30]. Such aggregates were not detected in the muscles of young WT and TG mice, and in old TG mice (Figure 7f, right panel), showing prevention of sarcopenia by ERRγ.

**Fig. 7.**
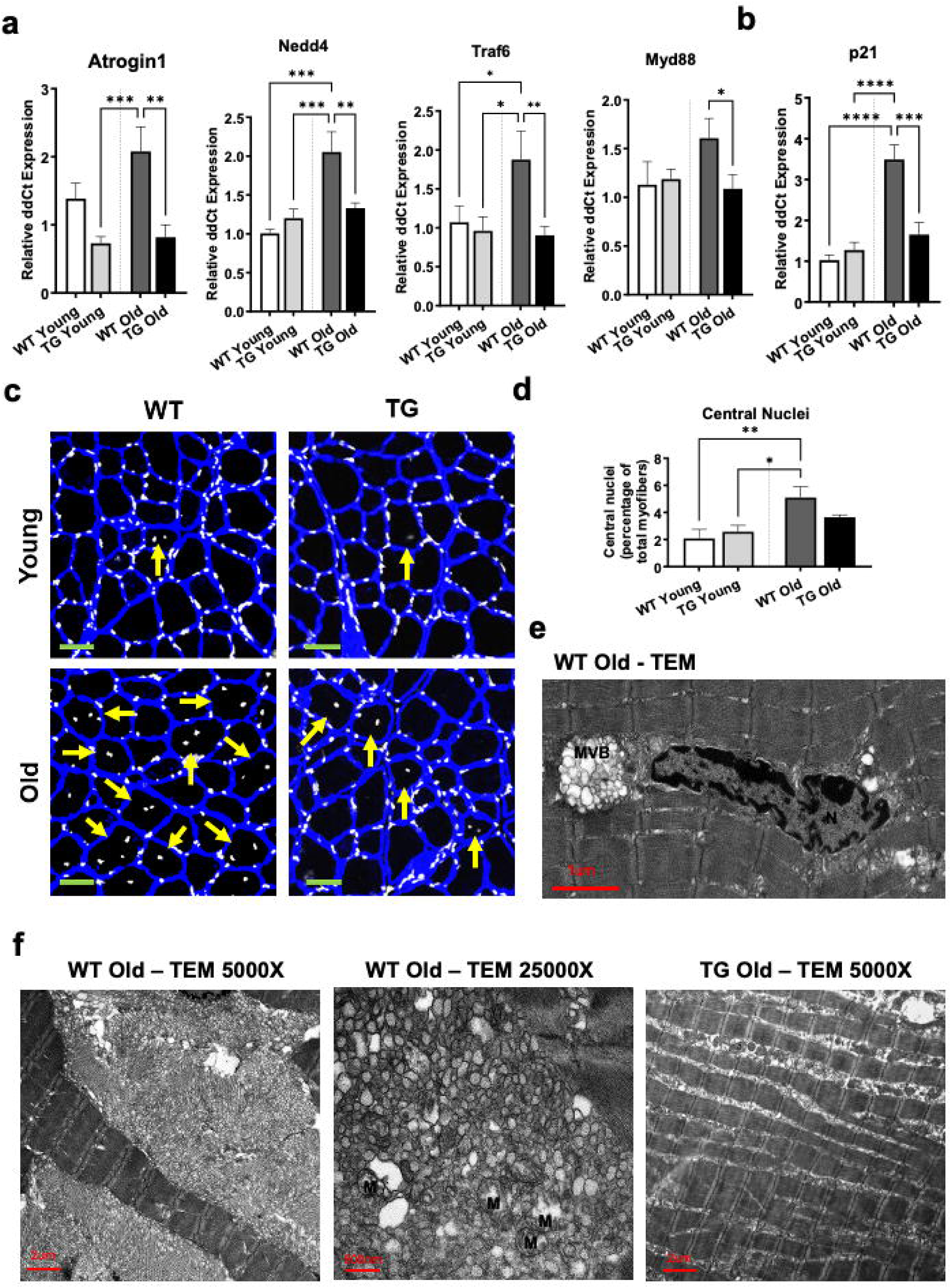
ERRγ suppresses hallmarks of atrophy and muscle aging in old mice. Following measurements were made in tibialis anterior (TA) or gastrocnemius from WT and TG mice at young and old age. **(a-b)** Quantification of atrophy (a) and senescence (b) gene in gastrocnemius. **(c)** Representative immunostained image of TA cryosections showing myofiber with central nuclei (yellow arrow). White – nuclei and blue – Laminin. Light green bar = 50 µm**. (d)** Quantification of centrally located nuclei (N=5). **(e)** Representative transmission electron microscopy (TEM) image of old WT TA showing centrally located nucleus along with multivesicular body (MVB). **(f)** Representative TEM micrographs of TA showing large aggregates with trapped mitochondria (labelled M in middle panel) in old WT TA (left and middle panel), which are absent in old TG TA (right panel). One-way ANOVA with Tukey’s post hoc test. p<0.05=*, p<0.01=**, p<0.001=***, and p<0.0001=****.

These data collectively indicate a dual effect of ERRγ in driving remodeling to damage-resistant oxidative myofibers, as well as preservation of type 2b cross-sectional area, along with preservation of vasculature and NMJ to combat hallmarks of sarcopenia.

### 3.6 Slow-twitch soleus is refractory to ERRγ-driven and age-dependent changes

Above analysis was performed in fast-twitch (TA) or mixed (gastrocnemius) muscles rich in sarcopenia prone type 2b myofibers. Soleus is pre-dominantly composed of slow-twitch type 1 myofibers. TG soleus has significantly higher gene expression of Esrrg but not Esrra compared to WT soleus from young and old mice (Figure S7a). However, the transgene did not significantly alter expression of representative genes for myosin isoforms (Figure S7b), mitochondria (Figure S7c), angiogenesis (Figure S7e), NMJ (Figure S7f), and atrophy (Figure S7g) in the young or old TG vs. WT mice. Therefore, slow oxidative muscles such as soleus are refractory to aging-dependent changes or ERRγ overexpression.

### 3.6 ERRγ preserves muscle contractile function and exercise capacity in old mice

Effect of ERRγ on muscle contractile function and exercise tolerance was measured in old and young mice. For contraction, we performed in vivo plantarflexion experiments by directly stimulating the muscle with subcutaneous electrodes. This enabled us to measure muscle contractile capacity by performing force-frequency analysis and energetics using fatigue protocol in conscious mice. Force-frequency analysis showed a significantly higher force produced in young WT vs. all other muscle groups at 60-140Hz, while the force production was comparable between rest of the groups (Figure 8a). The peak tetanic force produced in old WT muscle was lower than young WT muscle; whereas, the peak tetanic force production was comparable between old vs. young TG muscle (Figure 8b). The young WT muscle produced the highest force amongst the 4 groups. This corelates with the higher number and size of fast-glycolytic type 2b myofibers in young WT muscle compared to other groups. Contrastingly, in the TG mice type 2b myofibers have similar size, but are less in number compared to young WT mice resulting in lower basal peak force produced in transgenic muscle. After 3 minutes of rest following the force-frequency test, a fatigue protocol at sub-maximal frequency (50 Hz) was performed to measure the ability of the muscle to perform repeated contractions over 3 minutes. The total force produced over the first minute of the fatigue protocol is significantly lower in old vs young WT muscle (Figure 8c-d), indicating that the old WT muscle produces less force during repeated contractions. The old and young TG muscle produces comparable force throughout the fatigue protocol (Figure 8d). Maximum rates of contraction and relaxation are indicative of fatigue resistance, where faster rates corelate to less fatigue. The old WT muscle has the lowest maximal rates of contraction and relaxation, which is due to limited availability of ATP and compromises the ability of muscle to perform repeated contractions (Figure S6a-b). Both young and old TG muscle have faster maximal rates of contraction and relaxation than WT muscles (Figure S6a-b), and higher force in the latter half of the fatigue test (Figure 8c) indicating better muscle energetics.

**Fig. 8.**
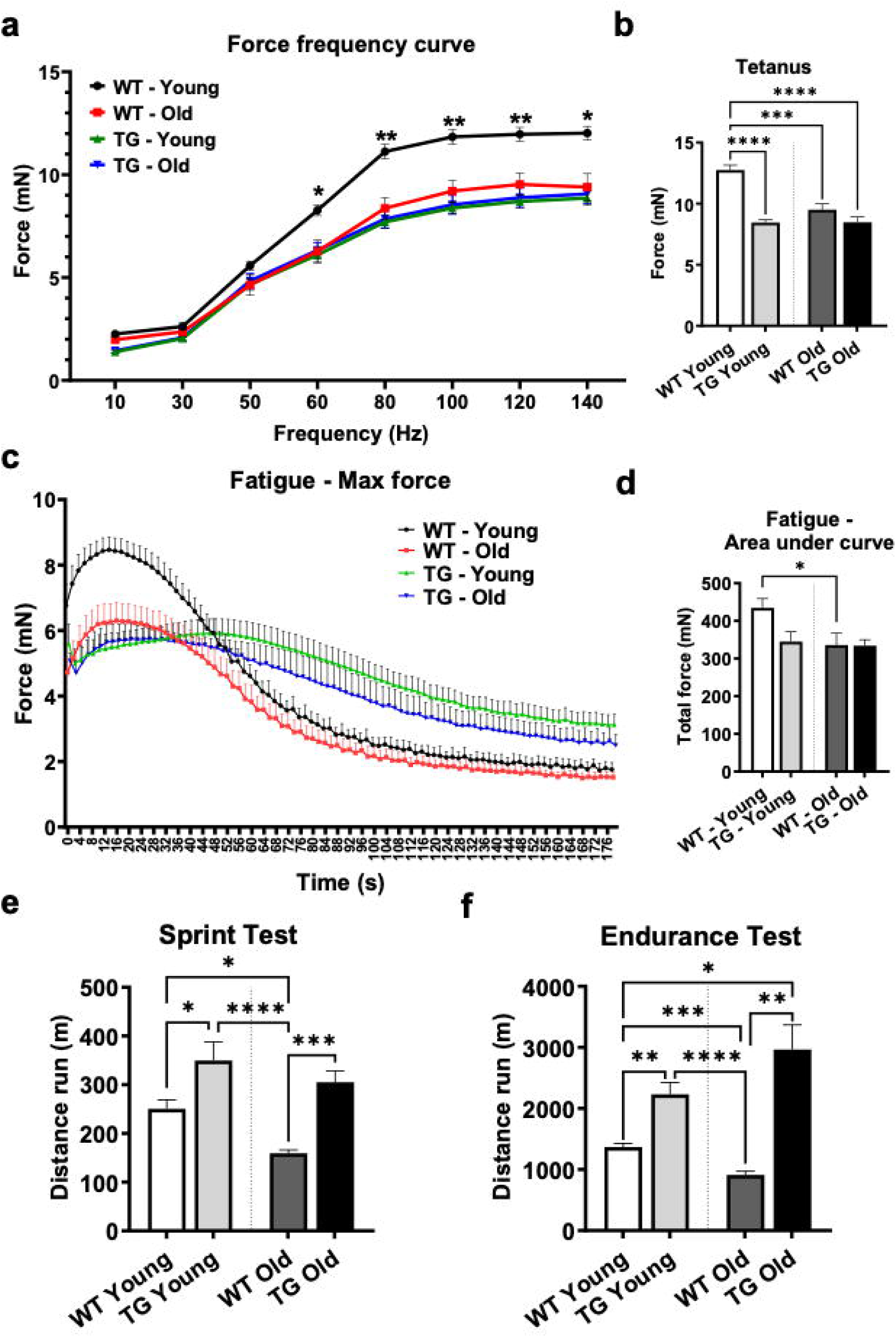
ERRγ preserves muscle function and exercise fitness with aging. Following measurements were made in WT and TG mice at young and old mice. **(a)** Force-frequency analysis of TG vs. WT mouse plantarflexion (N=5). Unpaired student’s t-test for comparison between young and old WT muscles. P<0.05=*, p<0.01=**. **(b)** Peak tetanic force production. **(c-d)** Graph showing peak force production every 2 seconds during a 3-minute fatigue test (c), and quantification of total force production during the fatigue protocol (d). **(e-f)** Quantification of treadmill running sprint (e) and endurance (f) test (N=9-12). One-way ANOVA with Tukey’s post hoc test. p<0.05=*, p<0.01=**, p<0.001=***, and p<0.0001=****.

The effect of ERRγ overexpression on exercise tolerance in young and old mice was measured using sprinting and endurance exercise treadmill test. The TG mice outperformed WT mice in the sprint test indicating a higher VO_2_ max (Figure 8e). Young TG mice also run longer distance in an endurance treadmill test compared to WT mice (Figure 8f). The old WT mice have lower exercise tolerance (sprinting and endurance) compared to young WT mice (Figure 8e-f, WT old vs. young). The old TG mice maintain their exercise tolerance with age (Figure 8e-f, TG old vs. young), outperforming the young mice indicating that ERRγ overexpression preserves exercise fitness in old mice.

## 4. DISCUSSION

We report that muscle ERRγ activation minimizes hallmarks of sarcopenia and maintains exercise fitness with aging. ERRγ drives oxidative switch, mitochondrial biogenesis, vascular and axonal/NMJ gene reprogramming that is beneficial in aged muscle. ERRγ prevents atrophy through oxidative myofiber type switch, preservation of type 2b myofiber size, and suppression of atrophy and senescence genes. ERRγ prevents age-related decline in muscle contractile function and exercise fitness. Therapeutic ERRγ activation could be a potential strategy for preventing or reversing sarcopenia.

Muscle ERRγ overexpression increases oxidative myofibers (type 2a and 2x), mitochondria and vascular supply, and exercise fitness in young mice [11]. Whether chronic ERRγ activation minimizes sarcopenia and age-related functional decline was unknown. We show that ERRγ transgene expression is sustained in old mice, along with its effect of increasing type 2a/2x myofibers, mitochondrial biogenesis, and vascularization in the muscle. RNA sequencing and validation revealed the underlying oxidative metabolic, mitochondrial and angiogenic transcriptional program age-independently driven by ERRγ transgene. TEM demonstrated a quantifiable increase in total mitochondrial area in TG vs. WT muscle in both young and old mice. There was a significant decrease in old vs. young WT total mitochondrial area in the intermyofibrillar regions, despite comparable mitochondrial gene/protein expression in old vs. young muscle. Nevertheless, several aerobic genes exhibited a downward expression with aging. The minimal suppression of angio/metabolic genes/proteins in old vs. young, despite clear deficits observed in TEM is likely due to mitochondria trapped within the large SR aggregates observed only in the old WT muscle. Our study does not detect an age-dependent decrease in angiogenesis and capillarity in muscle, as previously reported [31]. Nevertheless, ERRγ-induced mitochondrial and vascular program in old mice potentially contributes to preserving contractile function and exercise fitness.

We found that ERRγ is involved in NMJ formation. Several genes linked to NMJ biogenesis were upregulated by ERRγ in muscles of both young and old mice. Nrp1, Ptprm, and Pcdh17 are important for NMJ formation, axonal guidance, neurite outgrowth, and synapse formation. Aspa, gene encoding aspartoacylase is upregulated in glycolytic to oxidative fiber type switch and is protective in ALS induced muscle wasting [32]. Cxcr4 receptor activation promotes regeneration of motor axon terminals [33]. While NMJ numbers were decreased in the old muscle, ERRγ overexpression prevented age-related loss. ERRγ’s effect on mitochondrial homeostasis could contribute to NMJ regulation in muscle, as connection between mitochondria, redox balance, and NMJ formation is reported. Muscle-specific disruption of mitochondrial function and excessive ROS production through CHCHD10 or Drp1 deletion causes NMJ degeneration and denervation [34, 35]. Catalase-mediated restoration of muscle mitochondrial function and ROS reduction in SOD1 knockout mice (accelerated aging model) improves NMJ number and muscle innervation [36]. PGC1α and AMPK, regulators of muscle mitochondrial function and antioxidant effect, also regulate NMJ formation [37, 38].

ERRγ blocks age-dependent decline in total and average myofiber cross-sectional area. ERRγ prevented type 2b myofiber atrophy, preserving total and average type 2b myofiber cross-section area. Notably, total type 2b myofiber cross-section area and number was less in TG vs. WT muscles, due to ERRγ-dependent myofiber switch to oxidative type 2a/2x myofibers. Oxidative myofibers are less prone to sarcopenia compared to glycolytic myofibers [39] (also confirmed by observed resilience of soleus). Improved mitochondrial homeostasis by ERRγ could be involved in blocking myofiber atrophy. For example, inhibiting mitophagy worsens myofiber atrophy in muscle disuse [40]. Intramuscular dynamin like 1 protein (Drp1) overexpression prevents, whereas knockdown worsens muscle atrophy in aging mice [S1]. Parkin (another mitophagy regulator) overexpression also prevents age-dependent decline in muscle mass and strength. Muscle-specific deletion of OPA1 (mitochondrial fusion regulator) leads to mitochondrial deficiency and muscle wasting [S2]. Nrf2 deficiency (oxidative homeostasis) exacerbates age-dependent decline in mitochondrial dynamics and muscle mass [S3]. In addition to mitochondria, increased expression of angiokines, vascularity and perfusion in the aging skeletal muscle could retard muscle atrophy by driving muscle repair. Angiokine targets of ERRγ (Vegfa, Angiopoietin 1) are associated with muscle growth and repair [S4, S5]. Notably, proportion of myofibers with centralized nuclei (a marker of muscle damage) were elevated in old vs. young muscle but were absent in old muscles overexpressing ERRγ.

We did not detect age-dependent change in muscle ERRγ (or ERRα) expression. Other studies have shown decline in muscle ERRα/γ expression with aging across species, including humans [23, S6, S7]. Several regulators of ERRs transcriptional activity are repressed in aging. PGC1α and PGC1β, co-activators of ERRs, are downregulated in old vs. young muscle [S8]. AMPK activation, which is a stimulator of ERRs is impaired in the muscles of old vs. young mice [S9]. SIRT-1, a histone deacetylase, which interact with ERRs is downregulated with aging [S10]. Therefore, while muscle ERR expression remains intact with aging in our experiment conditions, their post-translational modifications, enrichment at target genes and/or transcriptional activity could be affected.

Our study has few limitations. ERRγ transgene is constitutively expressed in the skeletal muscle. Therefore, we observe the protective rather than therapeutic effect of ERRγ. Whether acute ERRγ activation in old mice elicits similar effects needs investigation. ERR pan-agonists that improve muscle fitness in young mice were recently reported [S11, S12], which could be tested in the aging model. Since our study was conducted in male mice, the effect of ERRγ on muscle aging in females remains to be tested. However, in ERRγ transgenic mice we did not find gender-based differences [11, 13]. As such, male mice show higher prevalence of sarcopenia than female mice [53]. ERRα was robustly increased by the ERRγ transgene. ERRs have overlapping muscle function [12, 14, 15]; therefore, the anti-aging effects of ERRγ could be partly mediated by ERRα. Our study does not decipher the relative contribution of each receptor to the observed effects.

These findings open avenues for future studies. ERRα/γ activation via gene therapy [S12, S14] or agonists [S11, S12] warrants inquiry as anti-sarcopenia therapy. The requirement of endogenous muscle ERRα/γ for healthy muscle aging should be studied in the muscle-specific ERRα/γ double knockout mice [12]. Interaction of ERRs with muscle growth (e.g. IGF1, Akt and mTOR) and/or atrophy pathways (e.g. myostatin, MuRF1, Atrogin1) needs exploration. Transcriptional landscape of ERRs, and their interactions with histones and co-regulators (e.g. PGCs, AMPK, Sirtuins) is not well-defined in aging, requiring in-depth genomic analysis. In summary, ERRγ has multifaceted muscle effects including mitochondrial biogenesis, muscle vascularization, oxidative myofiber switch, NMJ formation, and myofiber size preservation that maintains muscle quality, contractile function and exercise fitness with aging. ERRs could be potential targets for countering sarcopenia.

## ACKNOWLEDGEMENTS FUNDING SOURCE

This research was supported by National Heart, Lung and Blood Institute (R01HL152108), and Hamman Foundation Endowment to V.A.N; and American Heart Association grant to D.H.S. (23CDA1054599); and CCSG grant NIH P30CA016672. Z.Z. was supported by National Institutes of Health grants (R01LM012806 and U01AG079847). The RNA-seq data was generated in the UTHealth Cancer Genomics Core funded by the Cancer Prevention and Research Institute of Texas (CPRIT) grant (RP240610). C.C. is a CPRIT Postdoctoral Fellow in the Biomedical Informatics, Genomics and Translational Cancer Research Training Program (BIG-TCR) funded by CPRIT (RP210045). TEM was performed at the MD Anderson High Resolution Electron Microscopy Facility.

## ETHICAL STANDARDS

The manuscript does not contain clinical studies or patient data.

## DATA AVAILABILITY STATEMENT

The data that support the findings of this study are available on request from the corresponding author.

