## Supplemental Information for "Estrogen-related receptor signaling counters sarcopenia and preserves exercise fitness in naturally aged mice"

^1^Brown Foundation Institute of Molecular Medicine, McGovern Medical School, UTHealth, Houston, TX, 77030; ^2^MD Anderson UTHealth Graduate School of Biomedical Sciences, Houston, TX, 77030; ^3^Department of Biology and Biochemistry, College of Natural Sciences and Mathematics & ^4^Institute of Muscle Biology & Cachexia, College of Pharmacy, University of Houston. ^5^Center for Precision Medicine, McWilliams School of Biomedical Informatics, UTHealth, Houston, TX 77030

**^6^Corresponding Author:**

Vihang A. Narkar., Ph.D.

Brown Foundation Institute of Molecular Medicine

McGovern Medical School

UTHealth, Houston, TX 77030

**SUPPLEMENTAL METHODS**

**Animal husbandry**

We previously generated the muscle-specific ERRγ transgenic mice (TG) (1). These mice were housed in a temperature-controlled room (22°C) with ad libitum access to water and food (Pico Lab rodent diet 20; 13.2% fat) under 12:12 hours light-dark cycle. Up to 5 mice were housed in a single cage and age-matched young (4-5 months) and old (24-26 months) wild type (WT) and TG littermates were used. Animals were maintained and treated in accordance with the U.S. National Institute of Health Guide for Care and Use of Laboratory Animals. The procedures were approved by the Animal Welfare Committee at The University of Texas Health Science Center in Houston (UTHealth).

**Tissue collection**

Muscles were rapidly collected and flash frozen in liquid nitrogen or frozen in OCT on melting isopentane after the mice were euthanized by isoflurane inhalation followed by cervical dislocation.

**Quantitative real-time PCR**

mRNA was extracted from gastrocnemius and quantitative PCR (QPCR) was performed using SYBR Green PCR Master Mix (Applied Biosystems, Waltham, MA) with a BioRad XF96 cycler (BioRad, Hercules, CA). Data were normalized to Tbp.

**Western blotting**

Gastrocnemius was processed using RIPA buffer containing protease and phosphatase inhibitors using tissue lyzer and quantified using Pierce BCA protein assay kit, as previously described (2, 3). Samples were separated using standard SDS-PAGE and probed with 1:1000 dilution of primary antibodies for ERRα, ERRγ, CD31, PDGFRb, cytochrome C, and OXPHOS rodent antibody cocktail, as previously described (2, 3). Bands were visualized using chemiluminescence western blotting detection reagents.

**Immunohistology**

TA was cryosectioned at the mid-belly and consecutive sections were used for immunostaining. The sections were blocked in 10% goat serum in PBS. Myosin heavy chains 2a and 2b were stained using mouse monoclonal antibodies SC71 (5 μg/mL) and BF.F3 (4 μg/mL), respectively, and laminin using antibody L9393 (Sigma, St. Louis, MO) at 3.5 μg/mL dilution to visualize sarcolemma, as we previously described (2-4). CD31 antibody 10 μg/mL was used to detect capillaries. Motor neuron was stained using Neurofilament and Synapsin-1, acetylcholine receptor was stained using a-bungerotoxin. Alexa Fluor® secondary antibodies (Molecular Probes, Eugene, OR) were used to visualize all primary antibodies. Immunostained cryosections were examined using confocal microscopy. Myofiber number and area, CD31 number, and NMJ number were quantified using Image J.

**NADH-TR Activity Staining**

TA cryosections from young and old WT and TG mice were hydrated with PBS for 5 minutes followed by incubation with assay buffer (8mg/ml NADH and 10mg/ml Nitro blue tetrazolium in 0.05M Tris base pH7.6) for 15 mins at 37°C. The sectioned were washed with water and then unbound NBT was removed with three exchanges each of 30%, 60%, and 90% acetone in increasing and decreasing concentrations, followed by several rinses with water. The sections were mounted with aqueous mounting medium and imaged with brightfield microscope. Images were analyzed using Image J and data represented as densitometry of blue color.

**RNA sequencing and data analysis**

Total RNA was extracted with Purelink Kit (Ambion, Life technologies, Carlsbad, CA) and was further reverse-transcribed to cDNA with SuperScript III Reverse Transcriptase (Invitrogen, Waltham, MA) and used for RNA-sequencing analysis. Enriched poly (A) – tailed mRNA was used to prepare the RNA-sequencing library in the UT Cancer Genomics Center by following the instructions in KAPA mRNA HyperPrep Kit (KK8581, Roche, Holding AG, Switzerland) and KAPA Unique Dual-indexed Adapter kit (KK8727, Roche). RNA-sequencing with the 75 bp pair-ended running mode was performed in Illumina Nextseq550. Raw mRNA sequence reads were processed and bases with quality scores <20 and adapter sequences were removed with Cutadapt (v1.15) (5), followed by alignment of clean RNA-seq reads to GRCm38 with STAR (v2.5.3a) (6). Uniquely mapped reads overlapping genes were counted by HTseq-count with default parameter using annotation from GencodeM15. Only genes with >5 reads in at least one sample were retained. The raw read counts of retained genes were submitted for differential expression analysis of cases compared with controls with DESeq2 software, which uses a model based on the negative binomial distribution. the Benjamini and Hochberg’s approach was used to adjust the resulting P-values to control for the false discovery rate (FDR). Genes with fold change (FC) > 1.5 (or FC < 0.667) and FDR < 0.05 were assigned as differentially expressed genes (DEGs). Standard gene set enrichment analysis was performed with a hypergeometric test using WebGestalt (v 0.4.3). The resulting P-values were also adjusted using the Benjamini and Hochberg’s approach.

**Transmission electron microscopy**

Tibialis anterior muscle samples from young and old WT and TG mice were fixed with a solution containing 3% glutaraldehyde plus 2% paraformaldehyde in 0.1 M cacodylate buffer, pH 7.3, then washed in 0.1 M sodium cacodylate buffer and treated with 0.1% Millipore-filtered cacodylate buffered tannic acid, post fixed in1% cacodylate buffered osmium tetroxide + 1.5% potassium ferrocyanide and stained en bloc with 1% Millipore-filtered aqueous uranyl acetate. The samples were dehydrated in increasing concentrations of ethanol, infiltrated, and embedded in LX-112 medium. The samples were polymerized in a 60°C oven for approximately 3 days. Ultrathin sections were cut in a Leica UC Enuity ultramicrotome (Leica, Deerfield, IL), placed on formvar coated single slot copper grids, stained with uranyl acetate and lead citrate and examined in a JEM 1010 transmission electron microscope (JEOL, USA, Inc., Peabody, MA) at an accelerating voltage of 80 kV.  Digital images were obtained using AMT Imaging System (Advanced Microscopy Techniques Corp, Danvers, MA). 20-24 micrographs from 3 mice of each group were analyzed using ImageJ for mitochondrial number, average area, and total area per micrograph.

**In vivo muscle contraction**

The plantar flexion force measurements of the posterior right leg muscles were performed using 1300A 3‐in‐1 Whole Animal System (Aurora Scientific), as previously described (3). Peak tetanic force at 150Hz was recorded after which force-frequency data was obtained by stimulating the muscle for 400ms with frequencies of 10, 30, 50, 60, 80, 100, 120, and 150 Hz at 1-minute intervals. Following a 5-minute rest period, a fatigue protocol was performed where 400ms of a 50 Hz stimulation was given every two seconds for 3 minutes for a total of 90 stimuli. Contractile events were recorded with the ASI611A Dynamic Muscle Control software (Aurora Scientific) and were calculated with the accompanying ASI611A Dynamic Muscle Analysis software (Aurora Scientific).

**Treadmill test**

The maximal running speed of the mice was determined by treadmill sprint running test, as previously described (3, 7). Acclimatization of WT and TG mice to the treadmill running was performed over 3 days; on day 1 at a speed of 10 m/min for 10 min with 0° incline, 5° incline on day 2, and 10° incline on day 3. On day 5, the mice were placed on the treadmill and the speed, duration, and incline were changed as follows: 0 m/min, 5 min, 0° incline; 6 m/min, 5 min, 0° incline; 7, 8, 9, and 10 m/min, 30 sec each, 20° incline; and +1 m/min increment every min, 20° incline after that until exhaustion (5 sec off the treadmill). After 1 week of recovery, endurance test was performed, as described (3, 7). The speed, incline, and time were changed as follows: 6 m/min, 0° incline for 5 min; 6-14 m/min, 0° incline, +1 m/min increment every min; 14 m/min, 5° incline for 30 min; 16 m/min, 10° incline for 30 min; and 18 m/min, 15° incline until exhaustion. Exhaustion was defined as 10 sec off the treadmill on the platform, at which point the run was terminated.

**SUPPLEMETAL FIGURES**

**Supplemental Figure S1**

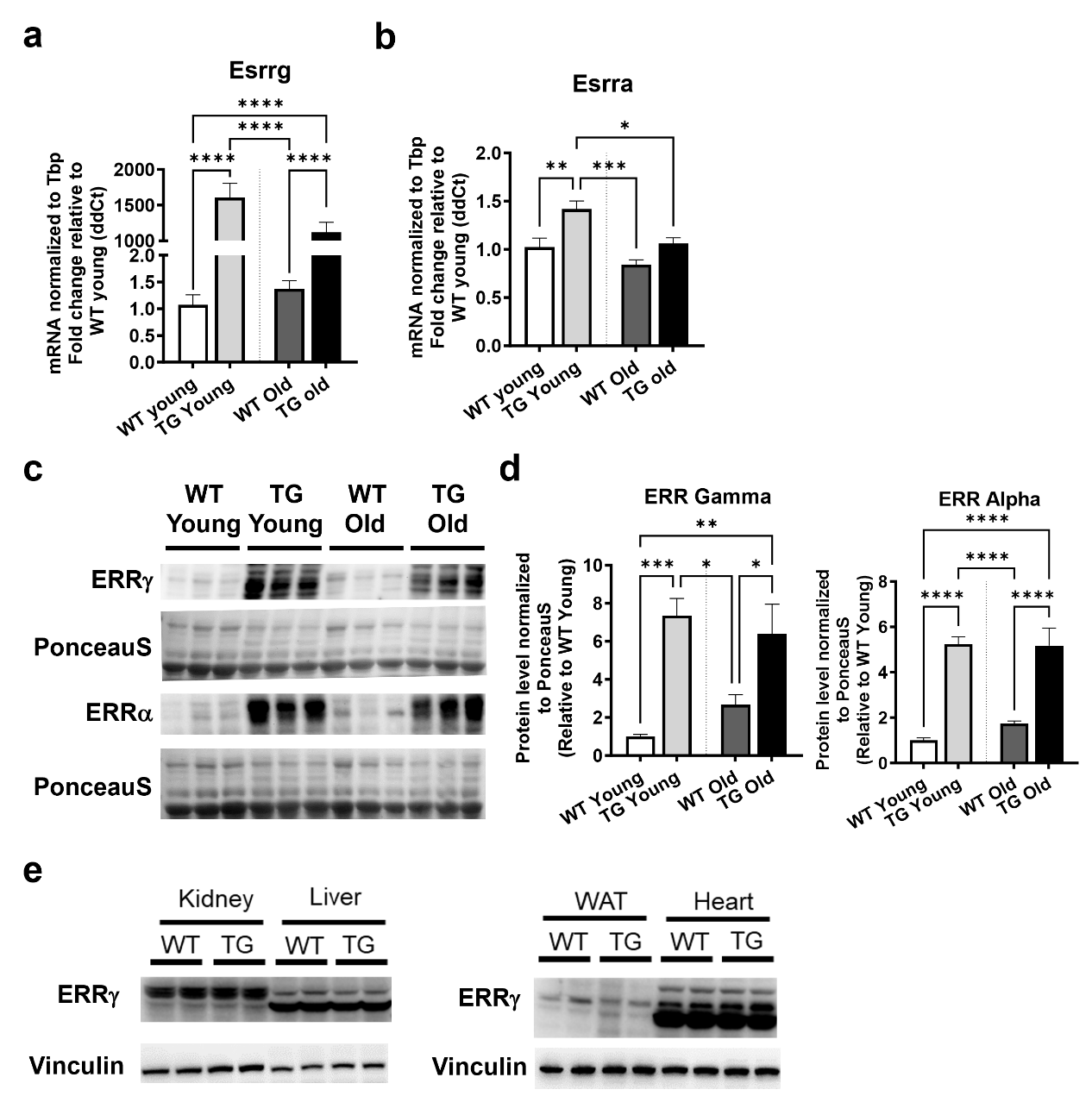

**Supplemental Figure S1. ERRα/γ expression in skeletal muscles of young and old mice.** Following was measured in wild type (WT) and transgenic (TG) gastrocnemius muscles of young and old mice (N=6). **(a-b)** Esrrg (a) and Esrra (b) gene expression. **(c-d)** Representative protein expression blot (c) and quantification (d) of ERRα and ERRγ. **(e)** Protein expression of ERRγ in non-muscle tissue including kidney, liver, white adipose tissue (WAT) and heart. One-way ANOVA with Tukey’s post hoc test. p<0.05=*, p<0.01=**, p<0.001=***, and p<0.0001=****.

**Supplementary Figure S2**

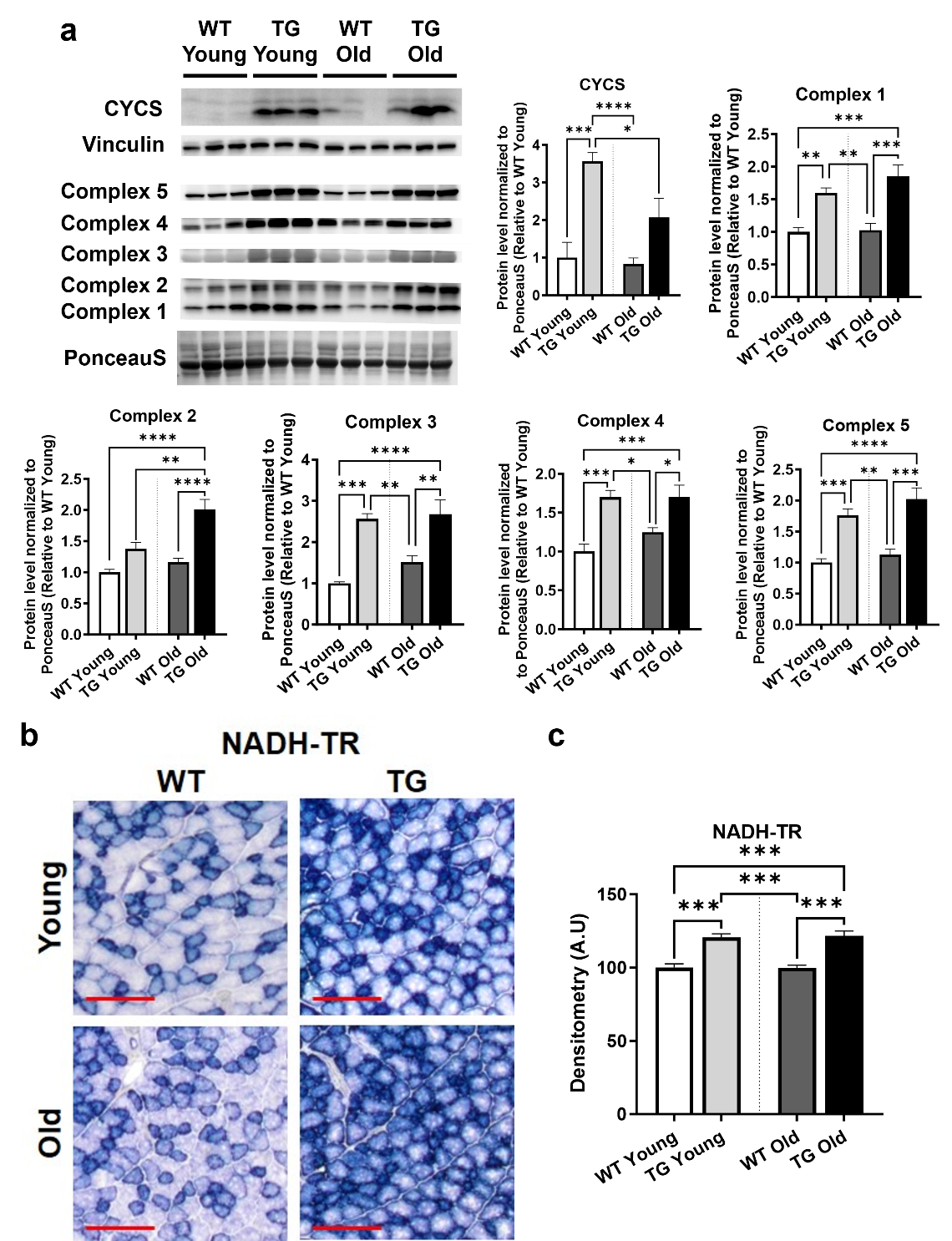

**Supplementary Figure S2. ERRγ induces mitochondrial biomarkers in skeletal muscles of young and old mice.** Following measurements were made in TG vs. WT muscles from young and old mice. **(a)** Representative western blot and quantification of CYCS and OxPhos subunits proteins in gastrocnemius (N=6). **(b-c)** Representative image of NADH-TR activity staining (b) and quantification (c) in TA cryosections (N=5 mice). Red bar = 200µm. One-way ANOVA with Tukey’s post hoc test. p<0.05=*, p<0.01=**, p<0.001=***, and p<0.0001=****.

**Supplementary Figure S3**

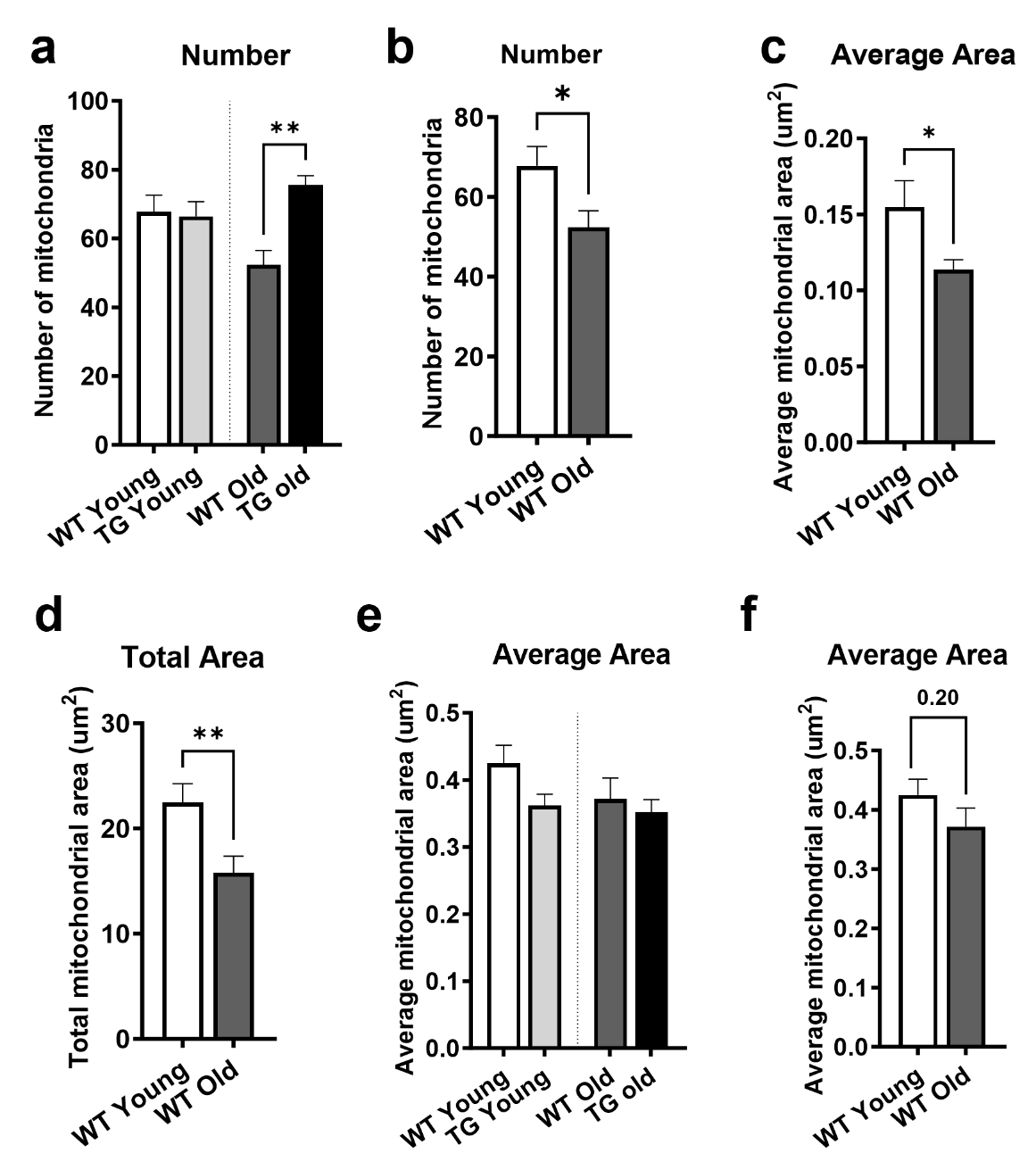

**Supplementary Figure S3. ERRγ increases mitochondrial content in TG muscle.** Following was measured in wild type (WT) and transgenic (TG) TA muscles of young and old mice (N=3). **(a)** Intermyofibrillar region shows significantly higher number of mitochondria in old TG vs. old WT muscle. **(b-c)** Quantification of intermyofibrillar region mitochondrial number (b), and average mitochondrial area (c) in old vs. young WT muscle. **(d)** Quantification of subsarcolemmal region total mitochondrial area in old vs. young WT muscle. **(e-f)** Quantification of subsarcolemmal region average mitochondrial area. N=20-24 micrographs from 3 mice in each group. One-way ANOVA with Tukey’s post hoc test. p<0.01=**. Unpaired Student’s t-test. p<0.05=*, and p<0.01=**.

**Supplementary Figure S4**

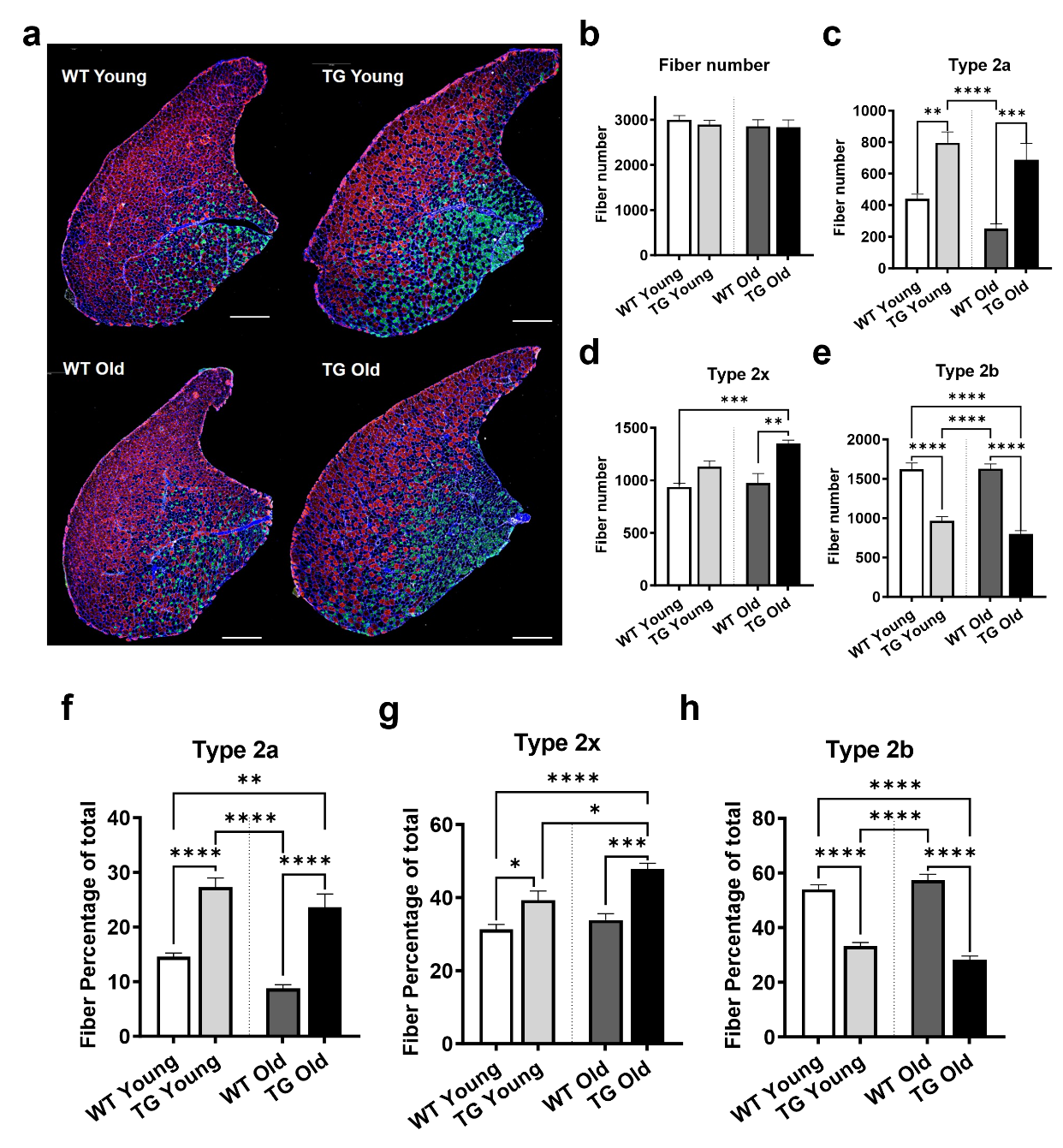

**Supplementary Figure S4. ERRγ drives oxidative myofiber type switch in young and old mice.** Following was measured in wild type (WT) and transgenic (TG) TA muscles of young and old mice (N=5). **(a)** Representative image of MHC immunostaining (Red – type 2B, Green – type 2A, Black – type 2X). White bar = 1000 µm. **(b-e)** Quantification of total (b), type 2a (c), type 2x (d), and type 2b myofiber number. **(f-h)** Quantification of percentage of type 2a (f), type 2x (g), and type 2b (h). One-way ANOVA with Tukey’s post hoc test. p<0.05=*, p<0.01=**, p<0.001=***, and p<0.0001=****.

**Supplementary Figure S5**

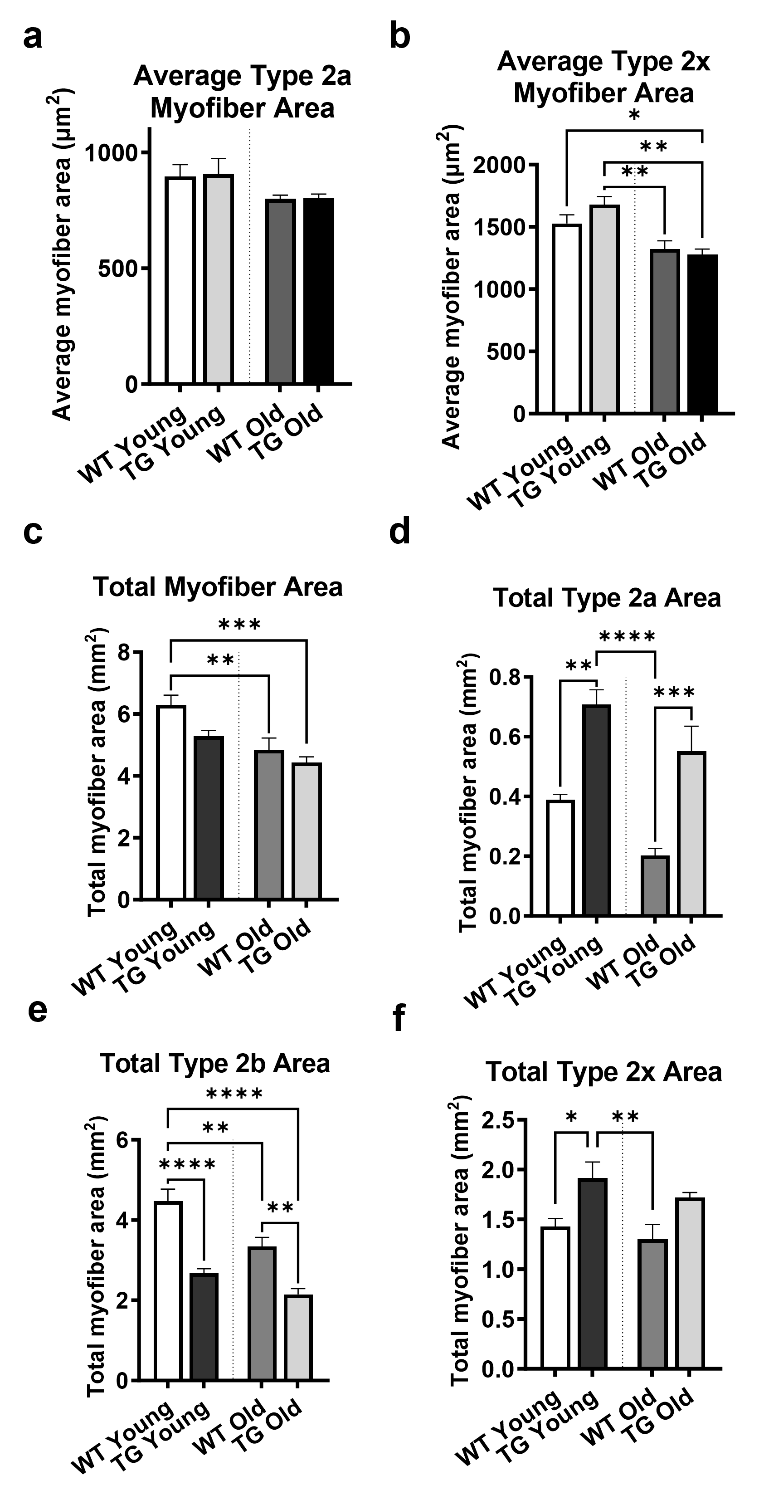

**Supplementary Figure S5. ERRγ preserves myofiber size in old mice.** Following was measured in wild type (WT) and transgenic (TG) TA muscles of young and old mice (N=5). **(a-b)** Average type 2a (a) and type 2x (b) myofiber area. **(c-d)** Total myofiber area (c), type 2a myofiber area (d), type 2b myofiber area (e), and type 2x myofiber area (f) in TA muscles of TG vs. WT mice at young and old age. One-way ANOVA with Tukey’s post hoc test. p<0.05=*, p<0.01=**, p<0.001=***, and p<0.0001=****.

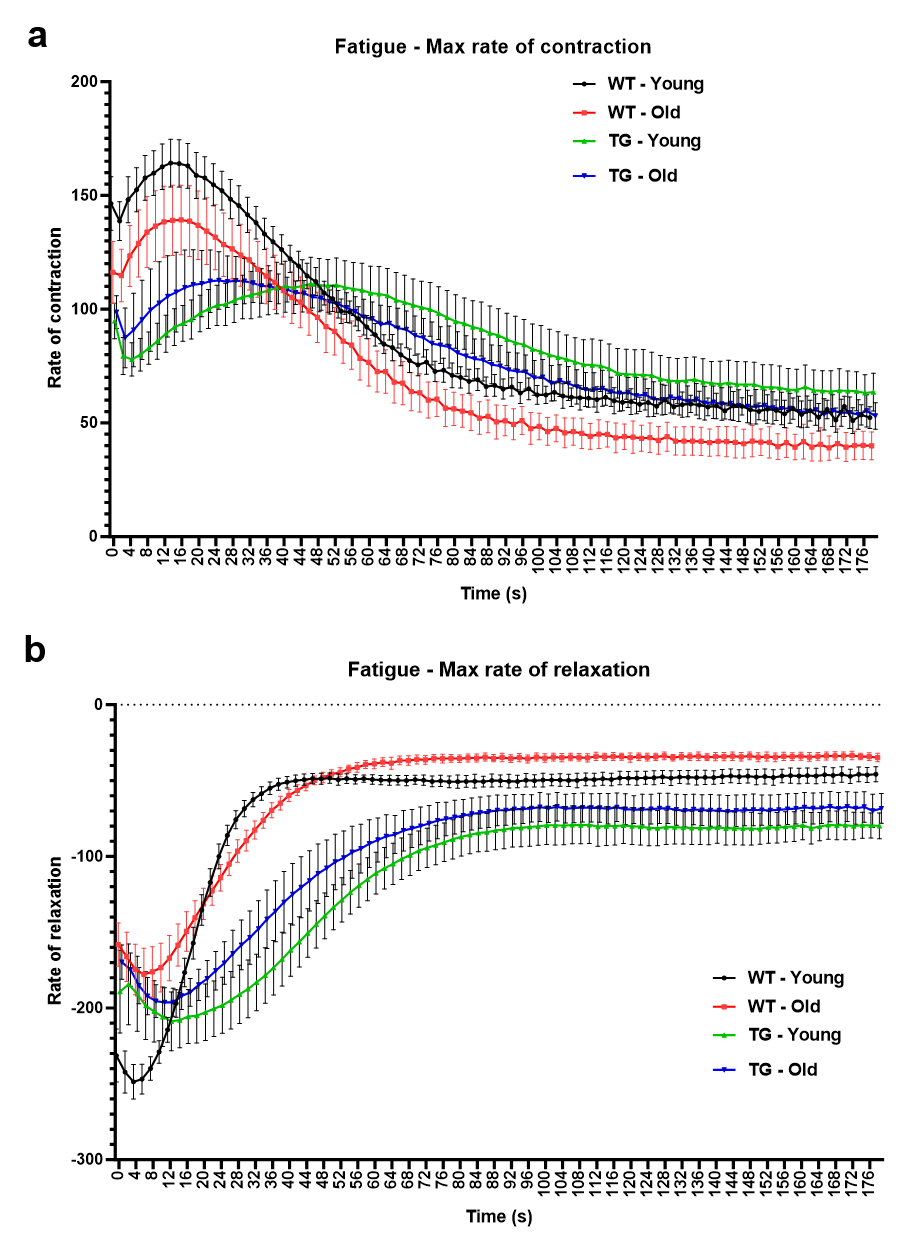
**Supplementary Figure S6**

**Supplementary Figure S6. Muscle contractile parameters in young and old mice.** Following was measured in wild type (WT) and transgenic (TG), young and old mice (N=6). **(a)** Graph showing the maximal rate of contraction at each time point during the in vivo plantarflexion 3-minute fatigue test**. (b)** Graph showing maximal rate of relaxation at each time point during the in vivo plantarflexion 3-minute fatigue test.

**Supplementary Figure S7**

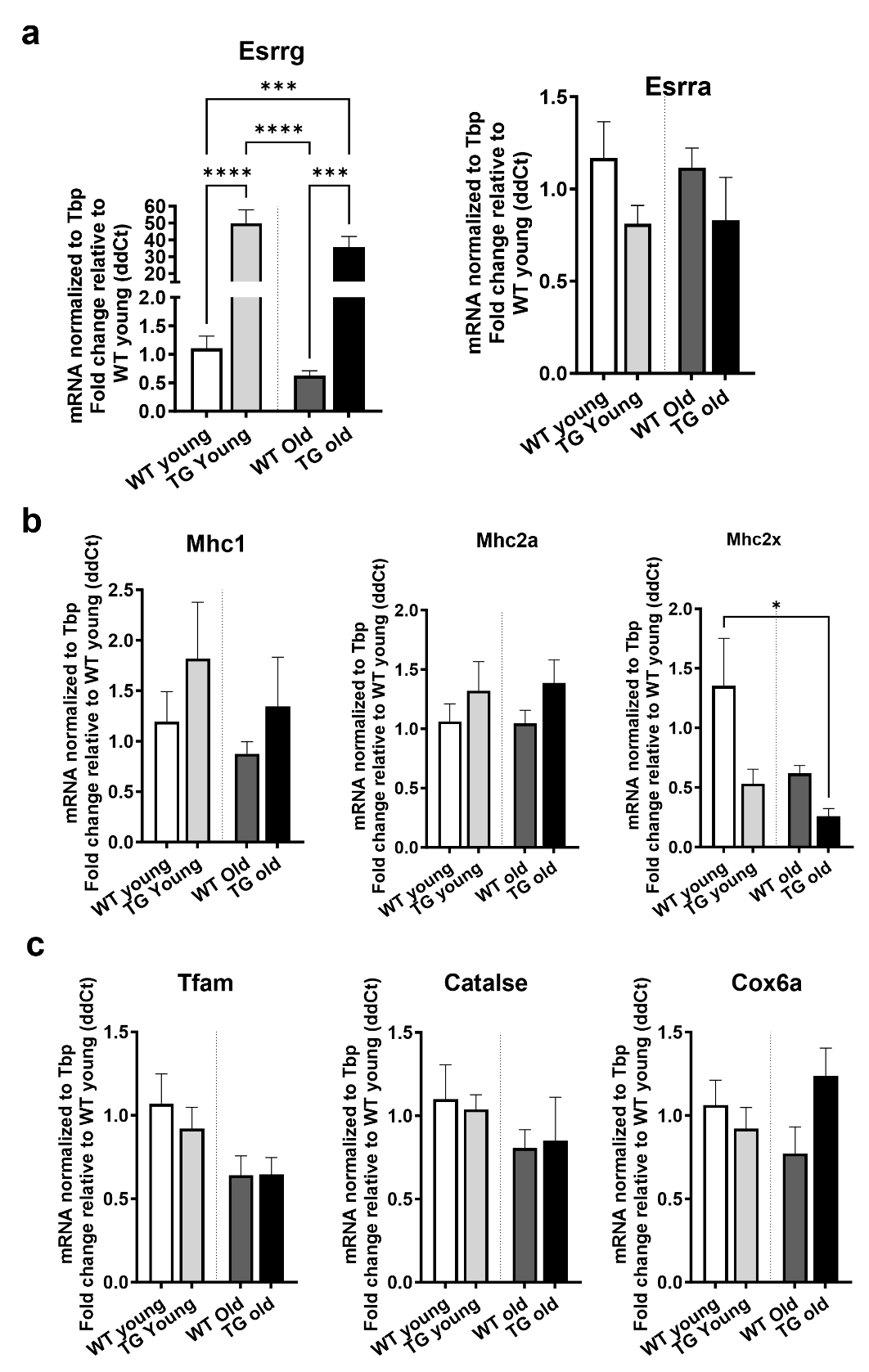

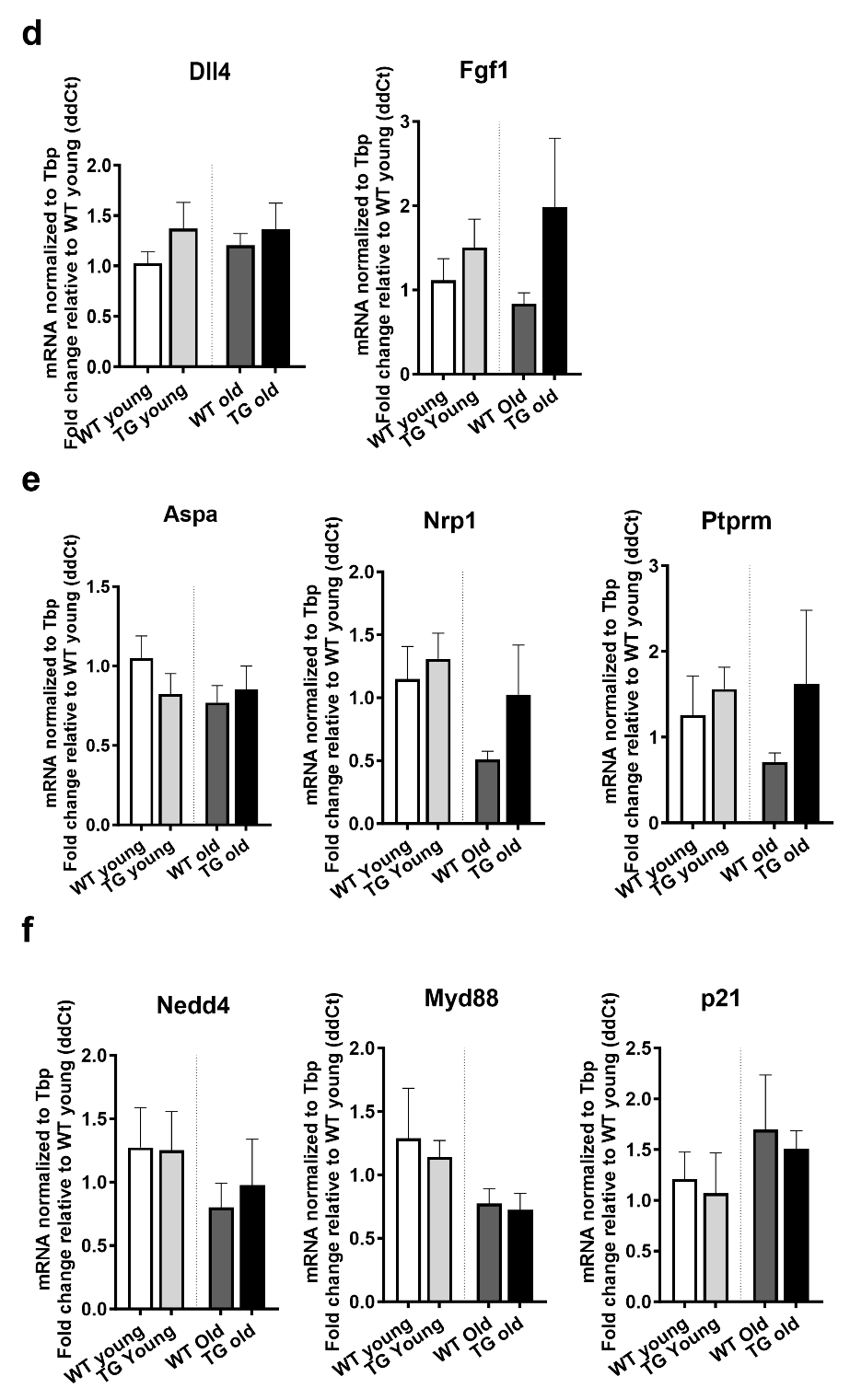

**Supplementary Figure S7. Oxidative slow-twitch soleus muscle is refractory to age-related changes and ERR**γ **overexpression.** Following was measured in wild type (WT) and transgenic (TG) soleus muscles of young and old mice (N=6). **(a)** Esrrg and Esrra gene expression. **(b)** Gene expression of myosin isoforms. **(c-f)** Expression of representative mitochondrial (c), angiogenesis (d), neuromuscular junction (e), and atrophy (f) genes. One-way ANOVA with Tukey’s post hoc test. p<0.05=*.
